# DOPAL initiates αSynuclein-mediated impaired proteostasis in neuronal projections leading to enhanced vulnerability in Parkinson’s disease

**DOI:** 10.1101/2021.06.15.448476

**Authors:** A. Masato, N. Plotegher, A. Thor, S. Adams, M. Sandre, S. Cogo, F. De Lazzari, C. M. Fontana, P. A. Martinez, R. Strong, A. Bellucci, M. Bisaglia, E. Greggio, L. Dalla Valle, D. Boassa, L. Bubacco

**Affiliations:** Department of Biology, University of Padova, Padova, Italy; Department of Neurosciences, University of California San Diego, La Jolla, California, USA; National Center for Microscopy and Imaging Research, University of California San Diego, La Jolla, California, USA; Department of Pharmacology, University of California San Diego, La Jolla, California, USA; Department of Neuroscience, University of Padova, Padova, Italy; Padua Neuroscience Center, University of Padova, Padova, Italy; Barshop Institute for Longevity and Aging Studies, University of Texas Health Science Center at San Antonio, Texas, USA; Department of Molecular and Translational Medicine, University of Brescia, Brescia, Italy

## Abstract

Dopamine dyshomeostasis has been acknowledged to be among the determinants of nigrostriatal neuron degeneration in Parkinson’s disease (PD). Several studies in experimental models and *postmortem* PD patients underlined increasing levels of the aldehydic dopamine metabolite 3,4-dihydroxyphenylacetaldehyde (DOPAL), which is highly reactive towards proteins. DOPAL has been shown to covalently modify the presynaptic protein αSynuclein (αSyn), whose misfolding and aggregation represent a major trait of PD pathology, triggering αSyn oligomerization in dopaminergic neurons. Here, we demonstrated that DOPAL elicits αSyn neuronal accumulation and hampers αSyn clearance at synapses and the soma. By combining cellular and *in vivo* models, we provided evidence that DOPAL-induced αSyn buildup lessens neuronal resilience, compromises synaptic integrity, and overwhelms protein quality control pathways, specifically at neuronal projections. The resulting progressive decline of neuronal homeostasis leads to dopaminergic neuron loss and motor impairment, corroborating the αSyn-DOPAL interplay as an early event in PD neurodegeneration.

## Introduction

Parkinson’s disease (PD) is an age-related severe degenerative syndrome with a multi-factorial pathology. Although different brain regions are affected by neurodegeneration depending on disease stage^1^, the highest degree of susceptibility is observed among the *Substantia Nigra pars compacta* (SNpc) dopaminergic neurons projecting to the *striatum*, which underlies the onset of cardinal motor phenotype^2^. According to the dying-back hypothesis for PD, synapse dysfunction and loss constitute the early pathological events initiating a progressive retrograde axonal injury, which gradually evolve to neuronal soma degeneration^3^.

The morphological, functional, and molecular features of SNpc dopaminergic neurons define the uniqueness of this neuronal subpopulation and its preferential vulnerability in PD^4, 5^. In addition to their autonomous pace making activity, dopaminergic neurons present complex arborizations of axonal projections, ensuring a profuse number of striatal synaptic connections. Importantly, the integrity of this neuronal network needs to be preserved through a high bioenergetic supply^6, 7^ and an efficient protein turn-over^8^.

Aging is among the prominent pathological factors leading to PD, as it represents the greatest challenge for upholding efficient degradative pathways^9^, thus altering neuronal protein quality control. Moreover, dopamine-induced oxidative stress seems to be paramount in nigrostriatal neuronal dysfunction as it can affect several intersected pathways, which lead to a negative loop of mitochondrial and lysosomal dysfunction, and protein aggregation^10^.

At striatal terminals, the misfolding and aggregation of αSynuclein (αSyn) has been identified as a driving factor in synaptic derangement^11–13^. αSyn is a small pre-synaptic protein known to regulate synaptic vesicles dynamics and exocytotic events^14, 15^. It is an intrinsically unfolded protein, which in part acquires an α-helical structure when associated with vesicle membranes^16^. Increased αSyn local concentration, oxidative stress and impaired neuronal proteostasis, promote αSyn conformational changes and aggregation, generating neurotoxic oligomers known to damage many neuronal pathways^17^.

In the last decades, the concept that a dyshomeostasis of catecholamines may lead to endotoxicity has been extended to dopamine catabolites, whose altered levels have been measured in autoptic samples and *in vivo* PD models^18^. Among them, the monoamine oxidize (MAO) dopamine catabolite 3,4-dihydroxyphenylacetaldehyde (DOPAL) is estimated to be by far the most reactive ^19, 20^. Although catechol oxidation to quinone species renders dopamine able to modify tiol groups, this is a spontaneous conversion with a slow kinetics when compared to the rate of enzymatic production of DOPAL by MAO, where the additional presence of the aldehyde moiety exacerbates DOPAL reactivity towards proteins^21^ with detrimental outcomes upon accumulation in the intracellular milieu.

The primary site of DOPAL burden is the pre-synaptic terminal, where its buildup is favored by a combination of defective dopamine storage in synaptic vesicles^22^, increased MAO activity with aging^23^, and decreased DOPAL detoxification by the aldehyde dehydrogenase enzymes (ALDH1A1, ALDH2)^20^. Accordingly, transcriptomic and proteomic studies in *post-mortem* PD patients brains, both familial and idiopathic, identified the selective reduced expression of ALDH1A1 among the molecular determinants involved in the preferential susceptibility of *SNpc* dopaminergic neurons^4, 24–26^, thus sustaining that the resulting DOPAL accumulation might be among the driving forces for dopaminergic neuron degeneration.

We previously demonstrated a functional consequence of DOPAL buildup at the pre-synaptic region, which induces a redistribution of synaptic vesicle pools in primary neuronal cultures^27^. This was linked to the generation of DOPAL-triggered annular-shaped αSyn off-pathway oligomers that were able to form pores on vesicles membrane. In agreement with other studies, we showed that DOPAL covalently modifies various lysines on αSyn sequence in a Schiff-base reaction between the primary amines and the aldehyde, with a higher and more specific reactivity than catecholamines^26–29^.

Here, we investigated the consequences of DOPAL buildup on neuronal homeostasis by triggering αSyn-mediated neurotoxicity. By combining biochemical studies with the advanced imaging technique of correlated light and electron microscopy (CLEM), we observed a DOPAL-induced αSyn accumulation among neuronal compartments and impaired αSyn clearance in primary neuronal cultures. We assessed the differential impact of the αSyn-DOPAL interplay in diverse neuronal districts, revealing altered synaptic integrity and overwhelmed degradative pathways in neuronal projections. Accordingly, these observations were substantiated in both mouse and zebrafish *in vivo* models of defective DOPAL detoxification, which exhibited altered proteostasis, dopaminergic neuron loss and impaired motor phenotype. Hence, our data disclose a novel pathological interplay between αSyn and DOPAL as a key molecular mechanism of enhanced vulnerability, which leads to synaptic dysfunction, compromised peripheral homeostasis and progressive dopaminergic neuron loss in the early events of PD.

## Results

### DOPAL triggers αSynuclein aggregation and affects αSynuclein proteostasis in neurons

Previous findings support the capability of oxidized dopamine to induce αSyn oligomerization by a non-covalent interaction of its catechol group with the _125_YEMPS_129_ motif at the C-terminus of αSyn^12, 30^. However, the additional presence of the aldehyde moiety on DOPAL reveals a diverse and higher reactivity that leads to a covalent and irreversible modification of multiple lysines on αSyn^27, 29, 31^. Using an *in vitro* αSyn aggregation assay following a time-course reaction (**Fig. 1a**), we observed substantial DOPAL-induced covalent modification of αSyn monomers detected by near Infra-Red Fluorescence (nIRF), and generation of SDS-resistant αSyn oligomers. These species displayed a completely different aggregation pattern as compared to that obtained upon incubation with dopamine, where the pool of monomeric αSyn remains unaltered.

**Figure 1.**
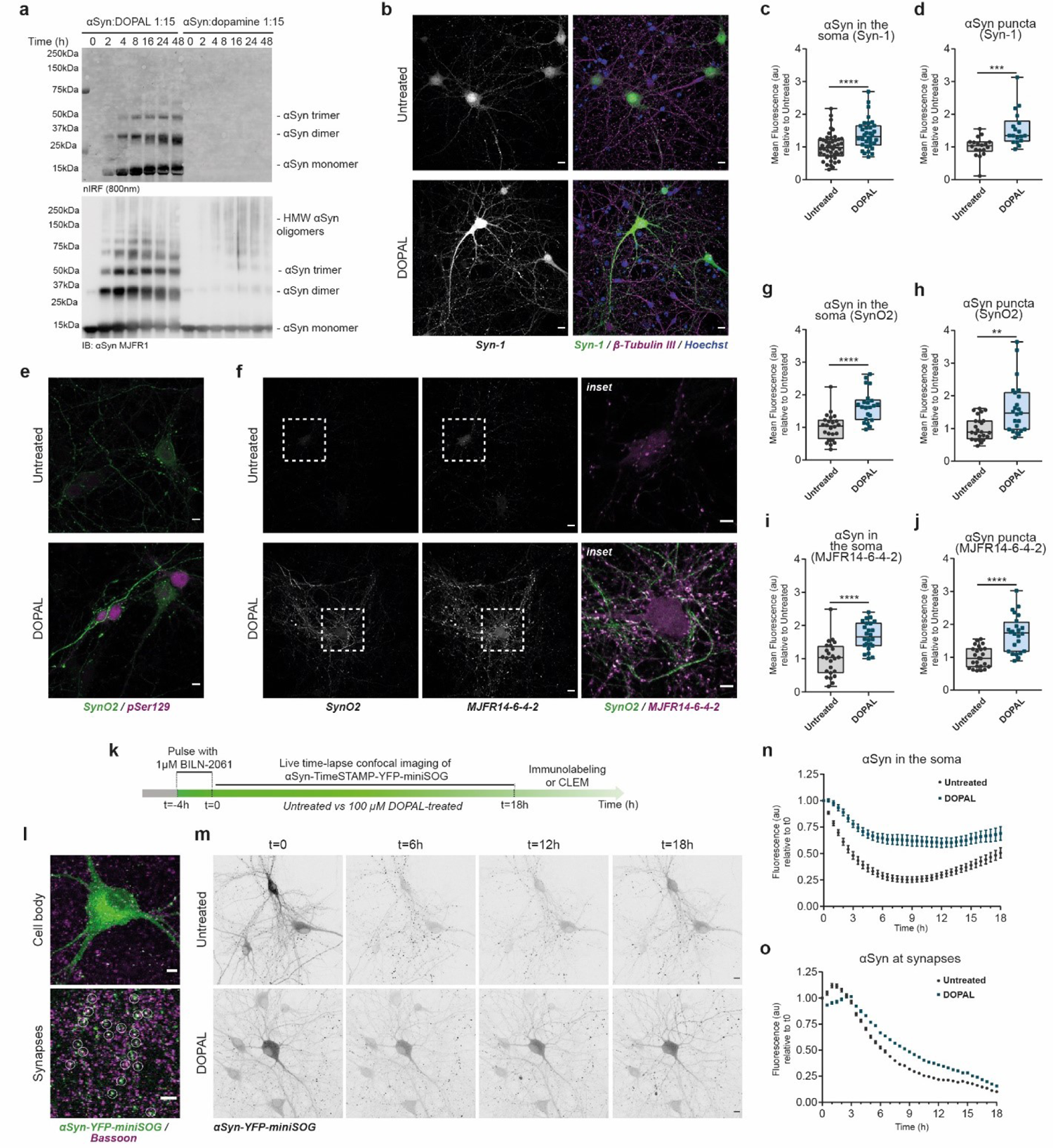
DOPAL affects αSynuclein proteostasis in primary neurons. (**a**) *In vitro* aggregation assay using 20 µM recombinant αSyn incubated with 300 µM (1:15) DOPAL or dopamine for different time-points, resolved by SDS-Page. The signal derived from oxidized DOPAL covalently bound to αSyn is acquired by nIRF at 800nm (top panel), whereas total αSyn oligomers are detected by immunoblot with the anti-αSyn MJFR1 antibody (bottom panel). (**b**) Immunostaining of αSyn (Syn-1) in untreated and 100 μM DOPAL-treated (for 24 hours) wild-type primary mouse cortical. In the right column, overlay of the αSyn signal (green) with the β-Tubulin III immunostaining (magenta). Nuclei are stained with Hoechst (blue). Scale bar: 10 μm. DOPAL-induced αSyn accumulation is expressed as (**c**) mean fluorescence intensity in neuronal soma and (**d**) mean fluorescence of αSyn-positive puncta. (**e**) Co-immunostaining of oligomeric (SynO2) and phosphorylated (pSer129) or (**f**) aggregated (MJFR14-6-4-2) αSyn in the same experimental conditions. Scale bar: 10 μm and 5 μm in the inset. DOPAL-induced aggregated αSyn buildup is expressed as (**g, i**) mean fluorescence intensity in neuronal soma and (**h, j**) mean fluorescence of αSyn-positive puncta. Data from (**c, d**) three and (**g, h, I, j**) two independent cultures are normalized to each untreated sample, pooled together, and analyzed by Mann-Whitney non-parametric test (** p<0.01, *** p<0.001, **** p<0.0001). (**k**) Schematic representation of the experimental setup of the pulse-chase experiment in αSyn-TimeSTAMP-YFP-miniSOG overexpressing primary rat cortical neurons. (**l**) Illustrative immunofluorescence images showing the αSyn-YFP-miniSOG localization in the cell body (green, upper image) and in the peripheral synapses, where it co-localizes with the pre-synaptic marker Bassoon (magenta, bottom image, white circles). Scale bar: 5 μm. (**m**) Snapshots at different time points of live time-lapse confocal imaging of rat primary cortical neurons expressing αSyn-TimeSTAMP-YFP-miniSOG, in untreated and 100 μM DOPAL-treated conditions. Scale bar: 10 μm. (**n-o**) Quantification of the YFP fluorescence variations in the soma and at synapses, where the fluorescence intensity of each cell body/synapse is normalized to t=0. Data are pooled together from three independent experiments (untreated: 41 cell bodies, 3320 puncta; DOPAL-treated: 38 cell bodies, 3712 puncta) and analyzed by Two-way ANOVA with Bonferroni’s multiple comparison test: (**l**) interaction **** p < 0.0001, treatment **** p < 0.0001; (**m**) interaction **** p < 0.0001, treatment **** p < 0.0001.

Considering the many potential neurotoxic outcomes associated with the accumulation of αSyn oligomers in the intracellular milieu, we set out to dissect the impact of DOPAL buildup on different processes that regulate αSyn proteostasis, including trafficking, subcellular localization, and clearance in the neuronal environment. To this aim, we employed diverse cellular models exposed to DOPAL treatment, including primary mouse and rat cortical neurons and the human neuroblastoma-derived BE(2)-M17 cells, to single out the effects elicited by DOPAL. Exogenously administered DOPAL was synthetized in our laboratory adapting Fellman’s protocol, which resulted in a compound with 95% purity (**Supp. Fig. 1**, see Online Methods for details on DOPAL synthesis and quality control). The detection of the nIRF signal derived from DOPAL adducts, present in both the detergent-soluble and insoluble fractions of BE(2)-M17 cells following an overnight treatment at increasing DOPAL concentrations, confirmed that DOPAL enters the cells (**Supp. Fig. 2a-b**).

**Figure 2.**
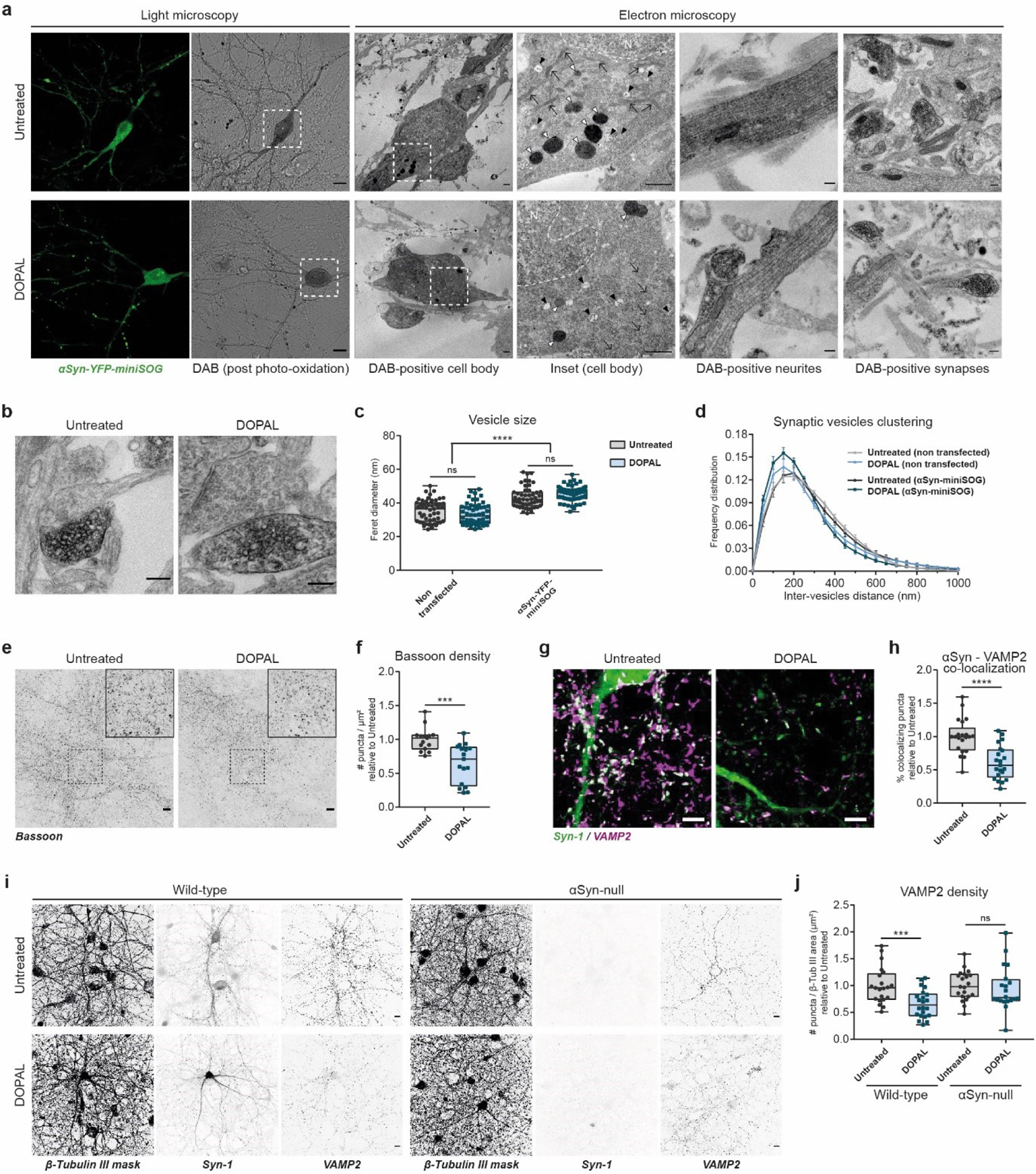
DOPAL-induced αSynuclein buildup affects synaptic integrity. (**a**) CLEM of rat primary cortical neurons expressing αSyn-TimeSTAMP-YFP-miniSOG, at t=24 hours after BILN-2061 pulse. Photo-oxidation in untreated and 100 μM DOPAL-treated neurons (scale bar: 10 μm) and representative electron micrographs of the DAB-positive cell bodies (scale bar: 1 μm), neurites and pre-synaptic terminals (scale bar: 200 nm). N: nucleus; ↗: mitochondria; △: MVBs; △: lysosomes. (**b**) Electron micrographs of pre-synaptic terminals in photo-oxidized areas showing αSyn-miniSOG-positive terminals in close proximity to terminals from non-transfected neurons. Scale bar: 200 nm. (**c**) Quantification of synaptic vesicles mean size measured as Feret diameter (nm) and (**d**) frequency distribution of synaptic vesicles clustering expressed as inter-vesicles distance (nm). Data are pooled from two independent experiments, three photo-oxidized areas (non-transfected _ untreated: 51 synapses; non-transfected _ DOPAL-treated: 54 synapses; αSyn-miniSOG-positive _ untreated: 49 synapses; αSyn-miniSOG-positive _ DOPAL-treated: 59 synapses), and analyzed by Two-way ANOVA with Sidak’s multiple comparison test: (**c**) **** p<0.0001); (**d**) interaction **** p<0.001, condition ns p>0.05. (**e**) Immunostaining of Bassoon in untreated and 100 μM DOPAL-treated (for 24 hours) wild-type primary rat cortical neurons (scale bar: 10 µm), with (**f**) the relative quantification of the density of Bassoon-positive puncta per area unit (µm^2^). (**g**) Immunostaining of αSyn (green) and VAMP2 (magenta) in untreated and 100 μM DOPAL-treated (for 24 hours) wild-type primary mouse cortical neurons (scale bar: 5 µm), with (**h**) the relative quantification of the percentage of αSyn-positive puncta colocalizing with VAMP2. (**i**) Immunostaining of β-Tubulin III (converted to binary mask) and VAMP2 in untreated and 100 μM DOPAL-treated (for 24 hours) wild-type and αSyn-null primary mouse cortical neurons (scale bar: 10 µm). (**j**) The synaptic density is expressed as number of VAMP2-positive puncta normalized to the β-Tubulin III area (µm^2^). Data from (**f**) three and (**h, j**) four independent cultures are normalized to each untreated sample, pooled together, and analyzed by Mann-Whitney non-parametric test (*** p<0.001, **** p<0.0001).

Furthermore, the specificity of the DOPAL-derived nIRF signal was confirmed by the quenching of DOPAL-mediated protein reactivity when co-treated with an excess of primary amines (i.e. aminoguanidine, AMG) in the cell medium (**Supp. Fig. 2c**).

On these premises, we next investigated the consequence of DOPAL exposure on αSyn steady-state levels in mouse primary neurons. Following a 24-hour treatment with 100 µM DOPAL, αSyn significantly increased in neuronal cell bodies as well as neurites and synapses (**Fig. 1b-d**, **Supp. Fig. 3a-b**), here identified as the scattered puncta detected in the field of view (see **Fig. 2g** for co-localization with the pre-synaptic marker VAMP2). αSyn elevation also correlated with an increased fraction of protein phosphorylation at serine 129 (pSer129), a recognized hallmark associated to αSyn pathological species, across different cellular models (**Fig. 1e**, **Supp. Fig. 3c-j**). Consistently, immunostaining with the anti-oligomeric αSyn (SynO2) and anti-aggregated αSyn (MJFR14-6-4-2) antibodies (which displayed a similar neuronal signal distribution as compared to the Syn-1 antibody staining) independently confirmed the DOPAL-induced accumulation of αSyn multimeric species both in the cell bodies and synaptic puncta (**Fig. 1f-j**, **Supp. Fig. 4**).

**Figure 3.**
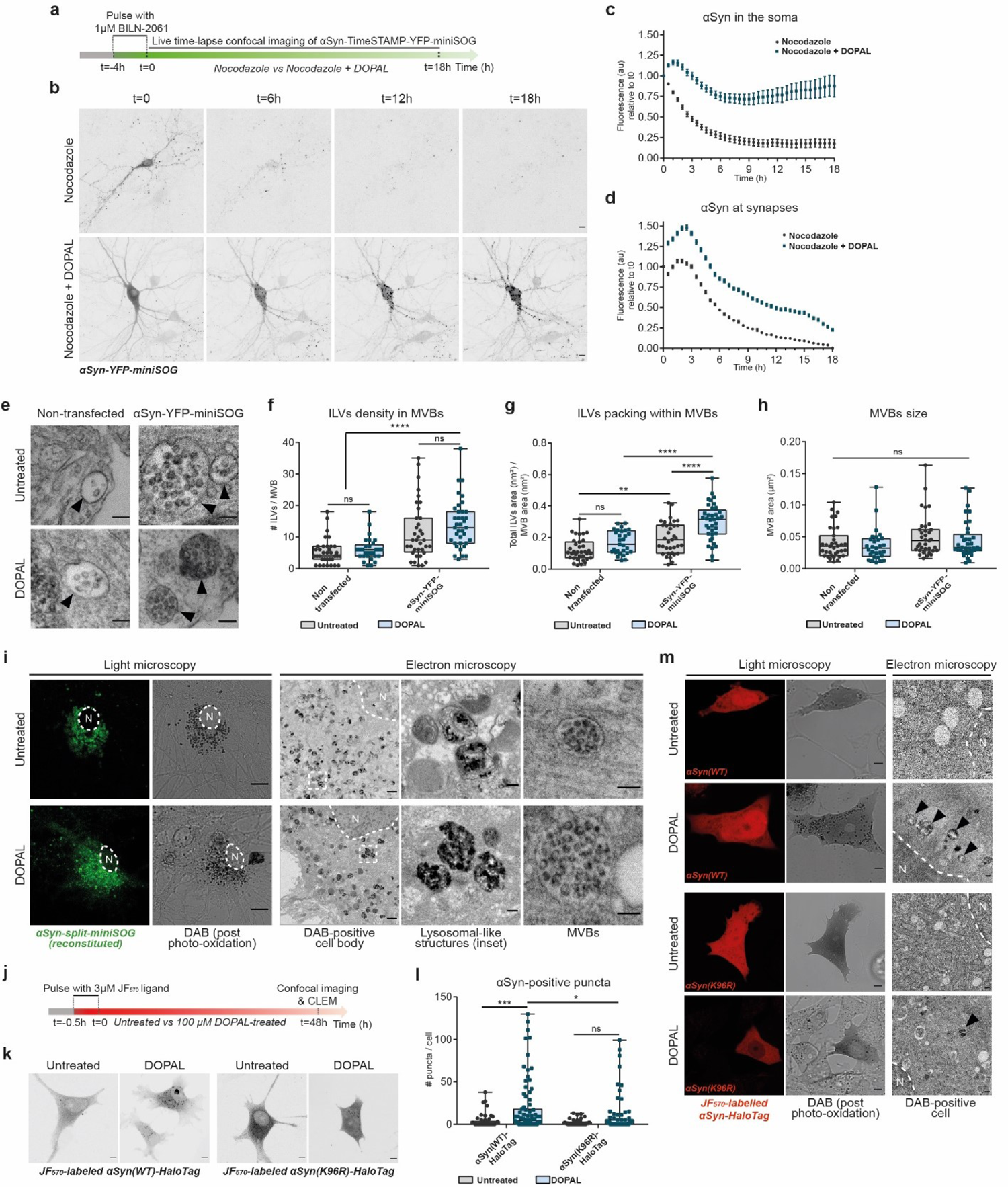
DOPAL promotes αSynuclein accumulation in the endo-lysosomal pathway. (**a**) Schematic representation of the pulse-chase experiment in αSyn-TimeSTAMP-YFP-miniSOG-overexpressing primary rat cortical neurons in the presence of 5 µg/ml Nocodazole in the cell medium, -/+ 100 µM DOPAL, and (**b**) representative snapshots at different time points. Scale bar: 10 μm. (**c-d**) Quantification of the αSyn fluorescence variations in the soma and at synapses, where the fluorescence intensity of each cell body/synapse is normalized to t=0. Data are pooled together from three independent experiments (Nocodazole: 32 cell bodies, 2154 puncta; Nocodazole + DOPAL: 23 cell bodies, 2600 puncta) and analyzed by Two-way ANOVA with Bonferroni’s multiple comparison test: (**c**) interaction **** p < 0.0001, treatment **** p < 0.0001; (**d**) interaction **** p < 0.0001, treatment **** p < 0.0001. (**e**) Representative electron micrographs of MVBs (indicated by the black arrowheads) in non-transfected and αSyn-miniSOG-positive neurons at t=24h after BILN-2061 pulse. Scale bar: 100 nm. Quantification of ILVs density in MVBs expressed as (**f**) number of ILVs per MVB and (**g**) total area of ILVs occupied in each MVB, and (**h**) MVBs area. Data are pooled from two independent experiments, three photo-oxidized areas (non-transfected _ untreated: 35 MVBs; non-transfected _ DOPAL-treated: 29 MVBs; αSyn-miniSOG-positive _ untreated: 36 MVBs; αSyn-miniSOG-positive _ DOPAL-treated: 37 MVBs) and analyzed by Two-way ANOVA with Sidak’s multiple comparison test (** p<0.01, **** p<0.0001). (**i**) CLEM of rat primary cortical neurons expressing reconstituted αSyn-split-miniSOG. Photo-oxidation of reconstituted αSyn-split-miniSOG in untreated and 100 μM DOPAL-treated neurons for 24 hours (scale bar: 10 μm) with corresponding electron micrographs of the DAB-positive cell body (N: nucleus; scale bar: 1μm) and higher magnification of perinuclear lysosomes and MVBs (scale bar: 100 nm). (**j**) Schematic representation of the pulse-chase experiment using the fluorescent JF_570_ HaloTag ligand in αSyn-HaloTag-overexpressing BE(2)-M17 cells. (**k**) Representative confocal images of BE(2)-M17 cells expressing αSyn(WT)-HaloTag and αSyn(K96R)-HaloTag, labelled with JF_570_ HaloTag ligand, in untreated and 100μM DOPAL-treated cells for 48 hours conditions. Scale bar: 5 μm. (**l**) Incidence of αSyn-positive cytoplasmic puncta. Data from three independent experiments are pooled together and analyzed by Two-way ANOVA with Sidak’s multiple comparison test (* p<0.05, *** p<0.001). (**m**) CLEM of BE(2)-M17 cells expressing αSyn(WT)-HaloTag and αSyn(K96R)-HaloTag, following protocol experiment as in (**j**). LM images (confocal and bright field, scale bar: 5 μm) and corresponding EM micrographs of DAB-positive cells (N: nucleus; scale bar: 200 nm) showing a prevalence of αSyn-positive lysosomal-like structures (green arrows) in DOPAL-treated αSyn(WT)-HaloTag-expressing cells.

**Figure 4.**
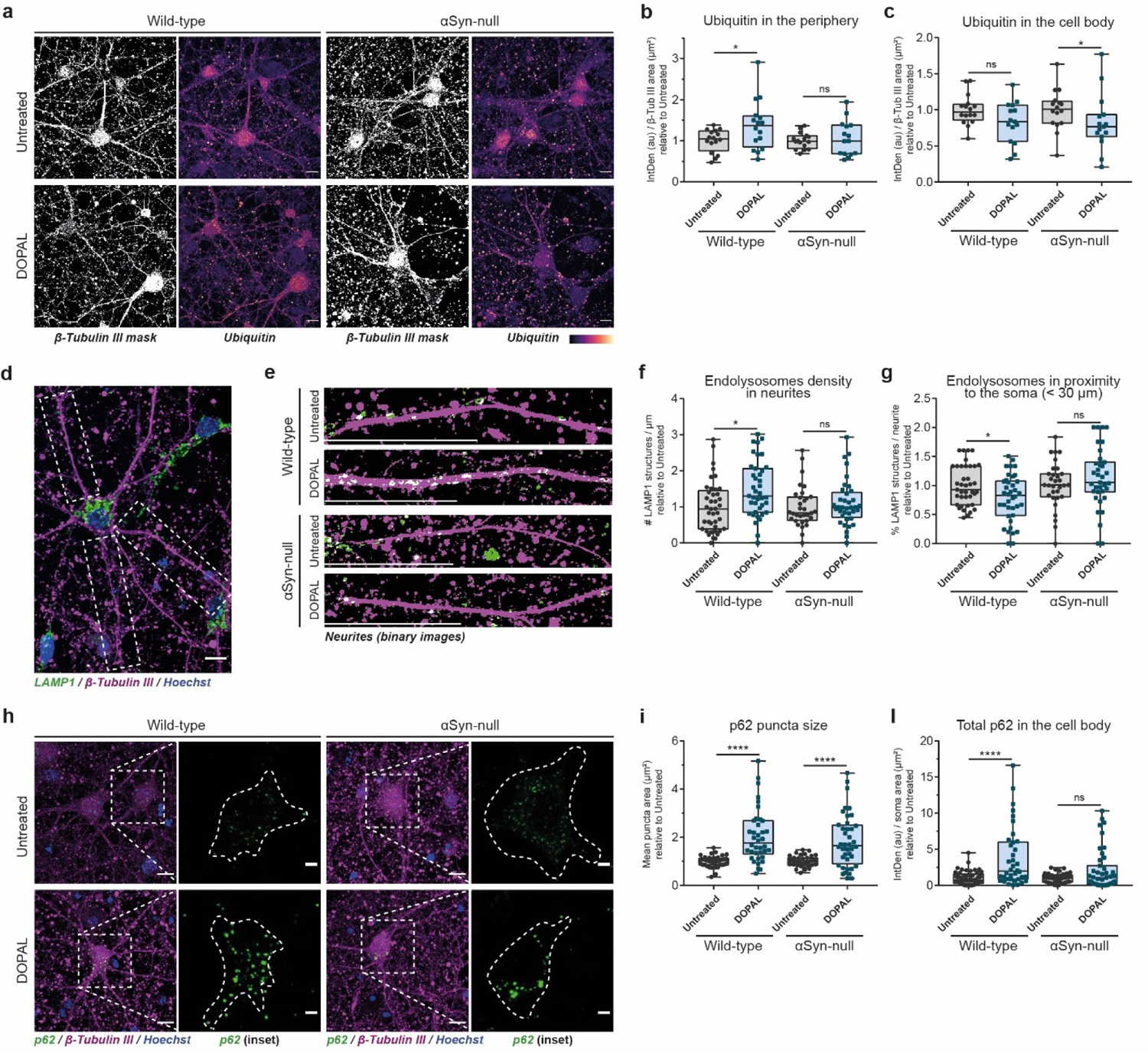
αSynuclein and DOPAL act in concert to hinder neuronal proteostasis. (**a**) Immunostaining of β-Tubulin III (converted to binary mask) and Ubiquitin (heatmap) in untreated and 100 μM DOPAL-treated (for 24 hours) wild-type and αSyn-null primary mouse cortical neurons. Scale bar: 10 µm. Quantification of the Ubiquitin fluorescence (**b**) in the periphery and (**c**) the cell bodies, both normalized to β-Tubulin III area (µm^2^). (**d**) Representative image of the immunostaining of β-Tubulin III (magenta) and LAMP1 (green) in primary mouse neurons. Nuclei are stained with Hoechst (blue). The insets provide examples of the criteria for neurite identification for endolysosomal density analysis in neuronal projections. Scale bar: 10 µm. (**e**) Enlargement of neurites analyzed in untreated and 100 μM DOPAL-treated (for 24 hours) wild-type and αSyn-null primary mouse cortical neurons. Both β-Tubulin III and LAMP1 fluorescence signals are converted to binary mask to emphasize the overlay (puncta in white). In the bottom left part of the images, the scale bar corresponds to 30 µm. (**f**) Endolysosomes density in neurites expressed as number of LAMP1 structures / µm^2^ and (**g**) percentage of endolysosomes in proximity to the soma (< 30 µm) in each neurite. (**h**) Immunostaining of β-Tubulin III (magenta) and p62 (green) in untreated and 100 μM DOPAL-treated (for 24 hours) wild-type and αSyn-null primary mouse cortical neurons. Nuclei are stained with Hoechst (blue). Scale bar: 10 µm. In the inset, the p62 fluorescence signal in the cell body is enlarged and the dotted line defines the soma boundaries (scale bar: 3 µm). (**i**) Quantification of the p62-positive puncta mean size (µm^2^) in each soma and (**l**) total p62 signal normalized to the soma area. (**b, c, f, g, i, l**) Data from three independent experiments are normalized to each untreated sample, pooled together, and analyzed by Mann-Whitney non-parametric test (* p<0.05, **** p<0.0001).

To gain insights into the mechanisms and consequences of DOPAL-modified αSyn accumulation, we next evaluated the kinetics of αSyn trafficking and subcellular distribution upon DOPAL treatment. We performed a pulse-chase experiment coupled to live-cell time-lapse confocal imaging by applying a CLEM approach using the αSyn-TimeSTAMP-YFP-miniSOG probe (Time – Specific Tag for the Age Measurement of Proteins)^32^ in primary rat neurons (**Fig. 1k**, **Supp. Fig. 5**). Briefly, the application of 1 µM BILN-2061 (here for 4 hours) induced the YFP-miniSOG fluorescent labeling of the newly produced αSyn, which progressively distributed within the soma where the protein is synthetized, and then along the neurites, and in the periphery, where it co-localized with the pre-synaptic scaffold protein Bassoon (**Fig. 1l**). Following BILN-2061 wash-out, live-cell time-lapse confocal imaging allowed to study the trafficking of the fluorescent αSyn subpopulation along the cell body and the synaptic boutons by analyzing the changes in the YFP-miniSOG signal over time (18-hour time-course) in the untreated condition or in the presence of 100 µM DOPAL (**Fig. 1m-o**). In the first 8 hours, untreated neurons exhibited a rapid 75% decrease of the fluorescence signal of αSyn-YFP-miniSOG in the soma, that was followed by a progressive rerise to 50% in the following 10 hours (**Fig. 1n**). In parallel, the fluorescence variations in the peripheral terminals indicated that, in these cells, the new fluorescent αSyn arriving from the soma quickly increased at the synapse showing a peak within the first 3 hours, which was followed by a progressive decrease (**Fig. 1o**). This trend is consistent with a combination of local clearance of αSyn both in the soma and the processes, together with a retrograde trafficking from the synapses^33^. Conversely, DOPAL-treated neurons exhibited a slower and less pronounced decrease of the fluorescence signal in the soma over time (**Fig. 1n**), and no increase in fluorescence within the first 3 hours at synaptic terminals, where the subsequent decrease of the protein was delayed as compared to untreated cells (**Fig. 1o**). These results indicate that DOPAL-treated neurons exhibit impaired αSyn trafficking as well as a less efficient degradation, as only 30% of the soma signal disappeared within the 18 hours analyzed.

**Figure 5.**
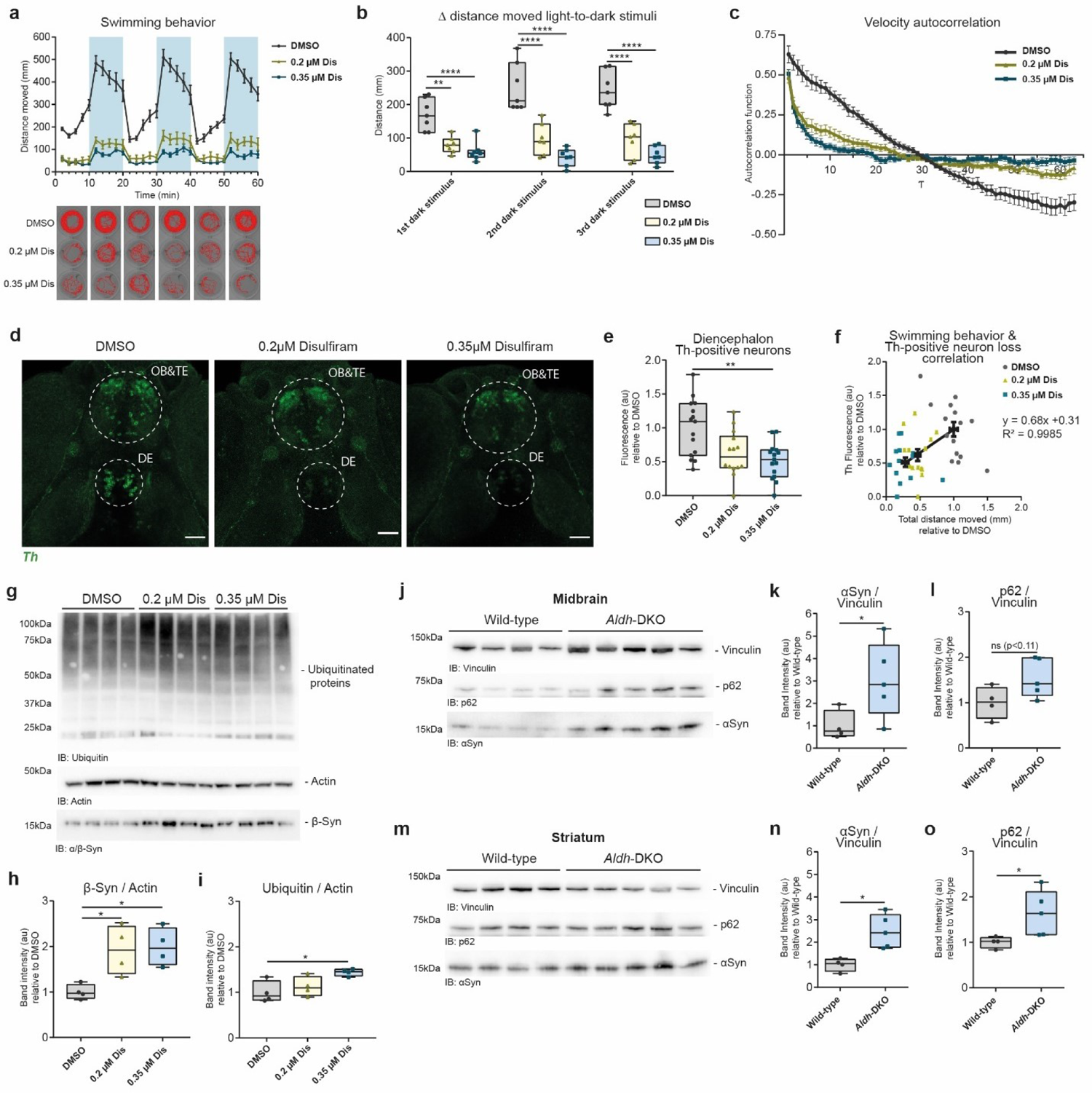
In *vivo* models of DOPAL neurotoxicity. (**a**) Time-course of the distance moved by DMSO-, 0.2 µM Disulfiram-(Dis) and 0.35 µM Disulfiram-treated zebrafish larvae during the light-dark locomotion test acquired at DanioVision™. In the graph, the blue areas indicate the three dark stimuli of 10 minutes each. Data are presented as mean ± SEM from seven independent experiments. In the bottom part, representative tracks for each 10-minute segment and each treatment are displayed. (**b**) Quantification of the delta distance moved in the first 2 minutes of each light-to-dark stimuli of the swimming behavior analysis. (**c**) Autocorrelation function of the time-course of velocity (10 second-bin) of movement during the light-dark test acquired at DanioVision™. Data are presented as mean ± SEM from three independent experiments (DMSO: 15 larvae; 0.2 µM Disulfiram: 14 larvae; 0.35 µM Disulfiram: 14 larvae). (**d**) Immunostaining of DMSO-, 0.2 µM Disulfiram and 0.35 µM Disulfiram-treated zebrafish larvae with the anti-TH antibody, which displays the dopaminergic neuron clusters of the olfactory bulb and telencephalon (OB&TE), and the diencephalon (DE). Scale bar: 50 µM. (**e**) Quantification of TH fluorescence signal in the dopaminergic neuron cluster in the diencephalon. (**f**) Correlation of swimming behavior (expressed as total distance moved during the light-dark test) of single larvae with the relative TH-fluorescence intensity in the diencephalon neuron cluster. Data are collected from three independent experiments (DMSO: 15 larvae; 0.2 µM Disulfiram: 14 larvae; 0.35 µM Disulfiram: 14 larvae) and both parameters are normalized to the mean value of the DMSO-treated sample. In the graph, the black dots represent the mean values (± SEM) for both parameters in each treatment, and corresponding equation of the linear regression fit. (**g**) Western blot of β-Syn and ubiquitinated proteins in lysates of zebrafish larvae heads, in DMSO-, 0.2 µM Disulfiram and 0.35 µM Disulfiram-treated conditions. Quantification of (**h**) β-Syn and (**i**) ubiquitinated proteins levels after normalization to Actin used as loading control. Data from (**b**) seven, (**e**) three and (**h,i**) four biological replicates are normalized to the mean value of the DMSO-treated samples and analyzed by Kruskall-Wallis non-parametric test with Dunn’s multiple comparison test (* p<0.05, ** p<0.01, *** p<0.001). Western blot of αSyn and p62 levels (**j-l**) in the midbrain and (**m-o**) in the *striatum* of 12-months old wild-type (n=4) and ALDH-DKO (n=5) mice. Band intensities are normalized to Vinculin as loading control. Data are normalized to the mean value of the wild-type for each read-out and analyzed by Mann-Whitney non-parametric test (* p<0.05).

### DOPAL-induced αSynuclein buildup affects synapse integrity

To understand the consequences of DOPAL buildup on αSyn subcellular localization and the organization of the neuronal ultrastructure, we exploited the miniSOG tag to perform CLEM studies in untreated and DOPAL-treated neurons at t=24 hours post BILN-2061 pulse-chase (**Fig. 1k**). The darker signal of the polymerized diaminobenzidine (DAB) imaged by EM reflected αSyn labeling and revealed its presence in the soma, in the neurites and in association with the membrane of synaptic vesicles (**Fig. 2a**), as previously described^34^.

We first studied the pre-synaptic terminals, where αSyn is thought to exert its physiological function^14, 15^. As αSyn has been reported to the dynamics and the membrane curvature of synaptic vesicles^11, 16^, we compared the size and clustering of vesicles in the synapses of non-transfected and αSyn-YFP-miniSOG expressing neurons (**Fig. 2b**). The measurement of synaptic vesicle size revealed an αSyn overexpression-driven increase in vesicle diameter (untreated neurons: non-transfected 35.3 ± 0.9 nm; αSyn-overexpressing 43.3 ± 0.9 nm). Interestingly, while DOPAL treatment (DOPAL-treated: non-transfected neurons 33.3 ± 0.9 nm; αSyn-overexpressing 45.4 ± 0.6 nm) did not significantly affect synaptic vesicle size (**Fig. 2c**), it did impact the synaptic vesicle clustering by shortening the inter-vesicle distance in an αSyn-dependent manner (**Fig. 2d**).

To further test the impact of DOPAL on pre-synaptic structures, we assessed synaptic density by immunofluorescence, detecting a considerably reduced number of Bassoon-positive puncta in primary rat neurons after 24-hour of 100 µM DOPAL treatment (**Fig. 2e-f**). Under the same treatment condition, a DOPAL-induced synaptic loss of comparable magnitude was independently confirmed by the analysis of the VAMP2-positive puncta density in primary mouse neurons. Also, the co-localization of αSyn peripheral puncta with this pre-synaptic marker showed a similar decrease (**Fig. 2g-j**). Notably, the DOPAL-induced synaptic loss relied on the presence of αSyn, as a non-significant reduction in VAMP2-positive puncta was observed in primary neurons isolated from mice with αSyn-null background (**Fig. 2i-j**). Overall, the data indicate that DOPAL-induced αSyn accumulation affects the synaptic integrity, altering vesicle organization and reducing synapse density.

### DOPAL affects αSynuclein turn-over and promotes αSynuclein accumulation in the endo-lysosomal pathway

Next, we sought to investigate the impact of DOPAL on αSyn turn-over, considering that *in vitro* DOPAL covalent modification of αSyn generates monomeric and oligomeric species that were more resistant to limited proteolysis by Proteinase K (PK) (**Supp. Fig. 6a-b**).

To assess whether DOPAL affects αSyn clearance also in the cellular environment, we performed pulse-chase live-cell imaging experiments in primary rat neurons and measured the fluorescence signal variations of αSyn-TimeSTAMP-YFP-miniSOG in the presence of Nocodazole^33^, which disrupts the microtubule network to exclude the confounding effect of the axonal trafficking component (**Fig. 3a**). Under these conditions, the impact of DOPAL on αSyn fluorescence decay was more pronounced (**Fig. 3b-d**). In the control neurons (not exposed to DOPAL), where the fluorescent αSyn was confined in the soma or in the peripheral terminals by Nocodazole, the protein was gradually degraded reaching 15% of the initial signal in 18 hours in the cell body, and it almost completely disappeared in the peripheral terminals. This allowed us to differentially estimate αSyn half-life in the range of 2.6 ± 0.4 hours in the soma and 6.3 ± 0.2 hours at the synapses, consistent with a rapid turnover of αSyn under the efficient neuronal protein quality control.

Conversely, in DOPAL-treated neurons, the decrease of αSyn fluorescence in the soma was considerably lower than in control cells, as the 85% of αSyn signal was still detected after 18 hours in the cell bodies (**Fig. 3c**). Meanwhile at the synapses, a rapid increase of the fluorescent signal in the first 3 hours was possibly due to the clustering of αSyn in DOPAL-induced aggregates, which appeared to be more resistant to proteolysis as their fluorescence decayed with a slower kinetics as compared to the untreated neurons, down to 20% of the starting protein signal at 18 hours (**Fig. 3d**).

Interestingly, in the CLEM experiments conducted in rat primary neurons using the αSyn-TimeSTAMP-YFP-miniSOG probe, we observed a consistent DAB-derived signal in Multi-Vesicular Bodies (MVBs) in the processes of both untreated and DOPAL-treated conditions, indicating the presence of αSyn in the lumen of Intra-Luminal Vesicles (ILVs) (**Fig. 3e**). By measuring ILVs density in MVBs (expressed either as number of ILVs per MVB or the fraction of area occupied by ILVs in the MVB lumen), αSyn overexpression appeared to increase ILVs loading as compared to MVBs of non-transfected cells, without affecting MVB size (**Fig. 3f-h**). Moreover, DOPAL treatment significantly increased the αSyn-positive ILVs packing within MVBs, when compared to MVBs in untreated and non-transfected neurons (**Fig. 3g**), thus highlighting an additive DOPAL effect on MVBs driven by the expression levels of αSyn.

Based on these observations of αSyn loading in MVBs, we hypothesized the engagement of the endo-lysosomal pathway in the clearance of αSyn aggregated species from the periphery to the soma. To investigate the distribution and accumulation of αSyn oligomers, we implemented a CLEM complementation assay using the αSyn-split-miniSOG probe^35^, which confirmed the labeling of αSyn oligomeric species in MVBs of DOPAL-treated neurons as well as in lysosomal-like compartments in the cell body (**Fig. 3i**). Accordingly, longer DOPAL treatment (100 µM DOPAL for 48 hours) in primary mouse neurons resulted in buildup of aggregated αSyn clusters in the soma and along the neurites in proximity to the cell body, detected by the MJFR14-6-4-2 antibody and quantified as decreased Nearest Neighbor Distance among clusters (**Supp. Fig. 6c-d**). Whereas, in untreated neurons, the αSyn staining was still consistent with the physiological distribution of the protein among the soma and the pre-synaptic terminals (more sparse puncta) as also observed for the αSyn staining with the Syn-1 antibody (**Supp. Fig. 6e**).

To further analyze the molecular mechanism behind DOPAL-induced αSyn loading in the endo-lysosomal pathway and clearance of αSyn oligomers via autophagy, we performed pulse-chase experiments using the HaloTag labeling technology to study αSyn turnover in the presence or absence of DOPAL, in a simpler paradigm. By using different HaloTag ligands to label αSyn-HaloTag over-expressed in BE(2)-M17 cells, we combined imaging (using the fluorescent JF_570_ ligand) and biochemical (using the biotin ligand) approaches to dissect the impact of DOPAL on αSyn degradation with both spatial and time resolution. Following a 30-minute pulse with 3 µM JF_570_ fluorescent HaloTag ligand to covalently label an αSyn-HaloTag subpopulation (**Fig. 3j**), 48-hour 100 µM DOPAL treatment led to the formation of αSyn-positive cytoplasmic puncta (**Fig. 3k-l**), which corresponded to αSyn accumulation in lysosomal-like compartments, as detected by CLEM (**Fig. 3m**). Moreover, knowing that the ubiquitination on lysine 96 by the ubiquitin-ligase Nedd4 is key in targeting αSyn to the endo-lysosomal degradation route^36^, we designed the αSyn(K96R)-HaloTag construct to prevent this specific post-translational modification on αSyn.

Interestingly, reduced αSyn(K96R)-HaloTag cytosolic inclusions and lysosomal accumulation were observed by CLEM in DOPAL-treated cells compared to αSyn(WT)-HaloTag (**Fig. 3k-m**), revealing compromised targeting of the mutant αSyn oligomers to the autophagic route, rather remaining diffused in the cytoplasm. This was substantiated by a complementary experiment in which, following an overnight 100 µM DOPAL treatment and a 3-hour pulse with 5 µM biotin ligand, αSyn(WT)-HaloTag and αSyn(K96R)-HaloTag monomeric and oligomeric species were isolated by pull-down with streptavidin-coated beads at different time points (**Supp. Fig. 7a**). In addition to a less efficient clearance of both αSyn(WT)-HaloTag and αSyn(K96R)-HaloTag in DOPAL-treated cells, the αSyn(K96R)-HaloTag oligomers showed a trend towards a higher accumulation compared to αSyn(WT)-HaloTag oligomers (**Supp. Fig. 7b-d**). Collectively, the data indicate that DOPAL affects αSyn turnover and K96 is required for the targeting to endo-lysosomal degradation.

### DOPAL and αSynuclein act in concert to hinder neuronal proteostasis in the periphery

Since we observed that DOPAL promotes αSyn oligomers to engulf the peripheral endosomal systems (**Fig. 3**), we then aimed at assessing whether an interplay between DOPAL-induced oligomeric αSyn burden and DOPAL-mediated protein modification could progressively affect neuronal proteostasis. We thus studied the levels and spatial distribution of different read-outs of degradative pathways in primary neurons following treatment with 100 µM DOPAL for 24 hours.

First, we evaluated whether DOPAL treatment in the presence (wild-type mouse neurons) or in the absence of αSyn (αSyn-null mouse neurons) equally affects protein ubiquitination, which drives protein quality control though ubiquitin proteasome system (UPS)-operated degradation as well as endosomal protein sorting and selective autophagy. Interestingly, we observed a significant DOPAL-induced accumulation of ubiquitinated proteins only in the periphery of wild-type neurons as opposed to the soma (**Fig. 4a-c**, **Supp. Fig. 8a**, see Online Methods for details on image analysis), supporting that αSyn synaptic accumulation is the key event driving the engulfment of quality control machineries following DOPAL buildup.

We then assessed whether the increased protein ubiquitination in the neuronal processes correlated with the upregulation of the endosomal pathway to transport aberrant proteins to the lysosomal degradation in the soma, as we observed for the MVBs by CLEM. Consistently, an increased density of LAMP1-positive endolysosomal structures was observed in the neurites of DOPAL-treated wild-type neurons when compared to the αSyn-null neurons (**Fig. 4d-f**), without significant difference in the detected organelle size (**Supp. Fig. 8b**). Of note, the endolysosomes in DOPAL-treated wild-type neurons accumulated mainly in the distal portion of the neurites (distance > 30 µm from the soma, **Fig. 4g**), supporting our hypothesis of impaired peripheral degradation systems by the DOPAL buildup-induced αSyn accumulation. Conversely, no significant difference in either organelle density or LAMP1-mean fluorescence intensity was observed in the soma endolysosomes, with the sole exception of a decreased in the average size of LAMP1-positive structures in αSyn-null neurons (**Supp. Fig. 8c-f**).

In addition to the endosomal pathway, more substrates are targeted to the lysosomal degradation via macroautophagy, a system involved in the clearance of misfolded aggregates, where the adaptor protein p62 binds ubiquitinated proteins and organelles to be engulfed in the autophagosomes that further fuse with the lysosomes. Interestingly, DOPAL treatment had a marked effect on p62 in the soma, generating larger and brighter puncta as compared to untreated neurons (independently from the genotype), synonymous of altered autophagic pathway (**Fig. 4h-i**, **Supp. Fig. 8g**). Although in wild-type neurons the p62 puncta density did not differ between untreated and DOPAL-treated neurons, in the αSyn-null cells a decreased number of p62 puncta was measured upon DOPAL-treatment (**Supp. Fig. 8h**), thus resulting in a significant increase in the overall p62 levels only when both DOPAL and αSyn accumulate (**Fig. 4l**). Collectively, these data show that DOPAL neurotoxicity strongly depends on its interaction with αSyn, whose oligomerization leads to impaired neuronal proteostasis.

### *In vivo* models of DOPAL neurotoxicity

According to the evidence presented so far, DOPAL buildup affects αSyn and the whole cellular proteostasis, representing a putative key molecular mechanism of enhanced neuronal vulnerability. Therefore, we aimed at recapitulating some of these aspects in diverse *in vivo* models of impaired DOPAL detoxification.

In a first *in vivo* model, we focused on the effect of the pharmacological inhibition of the Aldh enzyme in the brain of zebrafish larvae exposed to disulfiram (IC_50_ = 0.15 µM for hALDH1 and IC_50_ = 1.45 µM for hALDH2^37^), which hinders DOPAL detoxification and was reported to induce basal ganglia lesions after prolonged administration^38, 39^. Here, zebrafish larvae were administered 0.2-0.35 µM disulfiram in water from 2dpf (days-post-fertilization) to 6dpf. The disulfiram-exposed larvae displayed a significant impairment in the swimming behavior during the light-dark locomotion test (**Fig. 5a**). While the control larvae reacted to the dark stimulus by starting to swim faster in response to the sudden switching off the light, the treated larvae were unable to behave likewise, and they covered a significant reduced distance during the entire duration of the test (**Fig. 5b**, **Supp. Fig. 9a**). Moreover, the autocorrelation analysis of the velocity tracks during the light-dark locomotion test indicated that the disulfiram-exposed larvae had the tendency to display an intermittent swimming pattern compared to the control, suggesting a substantial impairment of the control of the voluntary movement (**Fig. 5c**, **Supp. Fig. 10**). We then asked whether the resulting impairment of the motor phenotype correlated with the loss of dopaminergic neurons of the diencephalon (DE) that is considered the nigral system of zebrafish^40, 41^. The treatment with disulfiram resulted in the selective dose-dependent loss of the Th-positive neuron cluster in the DE (**Fig. 5d-e**), while the Th-positive neurons of the olfactory bulb and the telencephalon (OB&TE) did not display a significant variation (**Supp. Fig. b**). More importantly, we were able to correlate the quantification of the swimming phenotype of the zebrafish individual larvae with their level of Th in the DE neuron cluster, thus confirming the specific neurotoxic effect due to the ALDH inhibition in the dopaminergic diencephalon neurons (**Fig. 5f**). Interestingly, along with the impaired swimming behavior and the dopaminergic neuron loss, we demonstrated that disulfiram exposure generated a significant accumulation of β-synuclein in the brain of exposed zebrafish larvae (**Fig. 5g-h**).

Although zebrafish lack of αSyn expression due to the putative loss of the ancestral *snca* locus during evolution^42^, the β-Synuclein isoform (*sncb*) displays a high sequence homology with human, mouse and rat αSyn at the N-terminus (where the epitope of the antibody used in immunoblot – aa 2-25 – maps) (**Supp. Fig. 9c**). Importantly, all the lysine residues in the αSyn sequence are conserved in the zebrafish β-Synuclein, therefore accessible to DOPAL modification. Also, in the same brain lysates increasing levels of ubiquitinated proteins were detected (**Fig. 5g,i**), supporting our hypothesis that DOPAL can lead to dysfunctional neuronal proteostasis *in vivo*.

We finally aimed at confirming our results in a genetic model of chronic endogenous accumulation of DOPAL. We analyzed midbrain and striatal brain tissues from *Aldh1a1^-/-^/Aldh2^-/-^* double knockout (*Aldh*-DKO) mice (**Supp. Fig. 9c-d**), which display an age-dependent parkinsonian-like phenotype and increased striatal DOPAL concentrations when compared to wild-type littermates^43^. Remarkably, we detected a significant increase in the levels of αSyn in both the midbrain and *striatum* (**Fig. 5j,k,m,n**) of 12 months old *Aldh1a1^-/-^/Aldh2^-/-^* mice with respect to wild type littermates. An accumulation of the autophagic cargo p62 was also measured, both in the midbrain and the *striatum* (**Fig. 5l,o**), independently supporting the paradigm of an altered proteostasis in the nigrostriatal neurons when DOPAL buildups.

## Discussion

In this study, we provided evidence on the impact of DOPAL-induced αSyn oligomerization on neuronal proteostasis, showing that DOPAL endotoxicity strongly depends on its interaction with αSyn and is exacerbated at synapses and neuronal projections.

Although the ability of DOPAL to covalently modify αSyn and trigger its oligomerization has been previously demonstrated^26–29^, we specifically unraveled a DOPAL effect on αSyn aggregation, localization, and clearance considering the complexity of neuronal compartmentalization. As we observed a significant increase in αSyn levels in all neuronal districts, the DOPAL-induced alterations in the live-cell time-lapse imaging experiments potentially derived from a combination of an impaired mobility of αSyn along the neurites and a decreased degradation efficiency both in the periphery and the soma, where different pathways are involved. At synaptic level, αSyn monomers and small *on-pathway* oligomers are known to be degraded by the UPS and local proteases, whereas in the cell body the autophagic pathway participate as additional backup, especially for the clearance of larger aggregates^44^. Interestingly, this might be reflected by the αSyn half-lives that we estimated in the two different regions in the live imaging with Nocodazole, where the αSyn turnover was about 2.5 times longer in the periphery as compared to the soma. Nevertheless, DOPAL-induced αSyn oligomers have been defined *off-pathway*, as they don’t acquire a unique structural conformation nor undergo fibrillation^27^. Moreover, the DOPAL modification on αSyn lysines is likely to interfere with the UPS-operated degradation. Hence, we speculated that the synaptic protein quality control machineries might not be fast enough to rapidly dispose the buildup of DOPAL-αSyn toxic species, thus requiring the engagement of the endosomal pathway to redirect these aggregates to the autophagic clearance in the soma. In support of this statement, in DOPAL-treated neurons, we observed an overload of oligomeric αSyn in MVBs, which are retrogradely transported to the soma where they fuse with the lysosomes to degrade their content. Interestingly, it has been recently demonstrated that neurons can deliver active lysosomes to distal axons to contribute to αSyn degradation from synapses, further supporting our hypothesis of the engagement of the autophagy-lysosomal pathway in the clearance of aberrant αSyn species in the neuronal projections^45^. Consistently, we observed a massive deposition of αSyn aggregates in lysosomal-like compartments in DOPAL-treated primary neurons and BE(2)-M17 cells by CLEM, suggesting a challenged autophagic pathway. Of note, the clogging of lysosomes and disruption of axonal network are reported to promote the fusion of the MVBs with the plasma membrane, releasing their vesicles in the extra-cellular space^46^. This is extremely relevant as exosomes represent a spreading pathway of αSyn toxic oligomers to neighbor neurons and glial cells^47^, thus augmenting both neurodegeneration and neuroinflammation.

We further proved that DOPAL buildup triggers αSyn-mediated neurotoxicity, compromising neuronal resilience. Interestingly, the greatest effects were observed at the neuronal distal regions and at the synaptic level, where αSyn concentration is higher. Here, αSyn reasonably represents a preferential target of DOPAL reactivity, as 10.7% of αSyn sequence is composed of lysine residues, a percentage which is higher than the average value of the lysine fraction in synaptic proteins (around 5%)^48^. At synapses, the DOPAL-induced αSyn accumulation appeared to affect synaptic vesicle organization and clustering, eventually reducing synapse density as reported by the VAMP2- and Bassoon-positive puncta quantification. Moreover, the distal regions are more vulnerable to the accumulation of misfolded and aggregated proteins, as they are less equipped with machineries for protein quality control and homeostasis^8^. A consistent pool of ubiquitin monomers is constantly available in the neuronal periphery, to ensure a rapid response in stress conditions and mediate the degradation of aberrant proteins by both the proteasome and the selective autophagy^49, 50^. Of note, we detected increased ubiquitin levels in the neuronal projections only in DOPAL-treated wild-type neurons, correlating with a higher density of LAMP1-positive structures in the distal portion of neurites and the overload of αSyn-positive MVBs. Both the synaptic injury and the overwhelmed endolysosomal pathway were observed to be dependent on αSyn expression, considering that the DOPAL-associated effects were limited in neurons with the αSyn-null background, thus corroborating the unique toxic interplay between DOPAL and αSyn.

On another note, it has to be considered that DOPAL presents a broad spectrum of reactivity and an impaired proteostasis induced by DOPAL modification would be more obvious on proteins with fast turn-over and prone to aggregation, as we observed for p62. In this case, a combination of DOPAL direct modification of p62 on lysines and cysteines (via the aldehyde and catechol moieties, respectively), p62 increased expression induced by antioxidant responses, and a blocked autophagic flux might all contribute to p62 accumulation and oligomerization^51^, which need further investigations.

As aggregated αSyn, highly phosphorylated at Ser129, ubiquitin, p62 and lipid membranes are the main constituents of Lewy Bodies (LBs)^52–54^, these observations become of interest for PD pathology. It may be speculated that, as αSyn-DOPAL oligomers burden impedes degradative pathways and neuronal proteostasis in general, this might lead to αSyn accumulation with enhanced *on-pathway* fibrillation, further contributing to LBs formation.

We finally tested our paradigm in *in vivo* models of DOPAL neurotoxicity, as a putative molecular mechanism of enhanced dopaminergic susceptibility of PD. Zebrafish larvae exposed to disulfiram, a potent ALDHs inhibitor, recapitulated a parkinsonian motor phenotype displaying decreased distance travelled during the light-dark locomotion test and a discontinuous pattern of movement. We also demonstrated that the impaired swimming behavior strongly correlated with the selective loss of Th-positive neurons in the diencephalon. Of note, the accumulation of Synuclein and the markers of degradative pathways were detected in both disulfiram-exposed zebrafish larvae and in aged *Aldh*-DKO mouse brain tissues. This evidence emerged as proof-of-concept of the impaired proteostasis when DOPAL detoxification is hampered, which reveals to be detrimental for the highly vulnerable dopaminergic neurons.

The proposed mechanism acquires relevance if one considers its specificity for dopaminergic nigrostriatal neurons, at the early stages of PD neurodegeneration. The translational relevance of our results also hinge on epidemiological studies that correlated an increased risk of developing PD with the exposure to drugs, pesticides, and chemicals that act as ALDHs inhibitors^20, 40^, enhanced by some genetic variations on *Aldh1a1* and *Aldh2* genes that were recently identified^20, 55, 56^. Unfortunately, only a few studies investigated the presence of DOPAL-modified αSyn species in PD autoptic samples, mostly because of the lack of reliable tools for their detection. Only one recent work reported increased levels of “αSyn dopaminylation” in the plasma of PD patients^57^. In this frame, the implementation of genetic screenings and suitable biomarkers oriented to the identification of subjects with increased risk of pathological DOPAL accumulation would be crucial for the early diagnosis of PD^58^. Such novel patient stratification strategies would also lead to a reconsideration of MAO inhibitors as therapeutic approach which, in past studies, did not match the requisites of disease-modifiers for PD.

## Acknowledgements

We thank Matthew Madany (UCSD) for providing the MatLab script for the image analysis of the live cell time-lapse imaging, Mason Mackey (UCSD) and Hiro Hakozaki (UCSD) for the training and assistance with the CLEM experiments, Junru Hu (UCSD) for the assistance with the primary rat neuronal cultures, Isabella Tessari (UNIPD) for the assistance in the generation of the αSyn-HaloTag constructs, the Zebrafish Facility (UNIPD) and Filippo Citton (UNIPD) for the zebrafish maintenance and assistance with the experiments. We thank Prof. Angelo Antonini (UNIPD) for the financial support of MS. RS is supported by grant number I01 BX001641 and Senior Research Career Scientist Award from the Department of Veterans Affairs Office of Research and Development. This work was financially supported by grants PRIN 2015 prot.2015T778JW (LB), PRIN 2017 prot.2017LYTE9M (LB, AB), Ing. Aldo Gini Foundation at UNIPD - Grant 2017 (AM), NIH R01GM086197 (DB), Branfman Family Foundation (DB), and NIH P41 GM103412 for support of the NCMIR directed by Prof. Mark Ellisman.

## Author contributions

AM designed research, performed the experiments, analyzed, and interpreted the data, and wrote the manuscript. DB and LB designed research, interpreted the data, and contributed to discussion and writing the manuscript. NP helped with the experiments, data analysis and interpretation, and manuscript editing. AT helped with the sample processing and imaging by TEM. SA synthetized the JF_570_ HaloTag ligand. MS and SC helped with the experiments, data interpretation and discussion. FDL and MB setup the protocol for DOPAL synthesis. CMF helped with the maintenance and experiments on zebrafish. PAM and RS generated the *Aldh*-DKO mouse model and provided the brain samples. AB provided contributed to the discussion. EG and LDV contributed to experimental design, data interpretation and discussion. All authors read and approved the final manuscript.

## Methods

### DOPAL synthesis

DOPAL was produced following the method by Fellman^59^, with slight modification. Briefly, 6 ml 85% ortho-phosphoric acid preheated to 120 °C in an oil bath were added to 400 mg of (±)-epinephrine hydrochloride (E4642, Sigma-Aldrich) in a 2-dram glass scintillation vial. The epinephrine-acid mixture was then vortexed until dissolved, re-submerged in the oil bath until it changed from yellow to a dark orange color and then added to 60 ml of H_2_O in a separating funnel. The water mixture was then extracted with 20 ml of ethyl acetate twice, washed with 10 ml of H_2_O and the combined organic layers were evaporated in a Savant SpeedVac concentrator until a constant mass was obtained. The resulting gel was first resuspended in 100% methanol, followed by evaporation by an N_2_ stream. DOPAL was then resuspended in distilled H_2_O and concentration was determined by assuming the same extinction molar coefficient of DA and L-DOPA (ε_280nm_ = 2.63 cm^-1^ mM^-1^). The quality of the DOPAL preparation was analyzed by reverse-phase HPLC (RP-HPLC) on a Phenomenex Jupiter column (300 Å/5 μm, 250 mm × 4.6 mm), using linear gradients of solvent B in eluent A (A: 0.1% TFA in H_2_O, B: 0.08% TFA in acetonitrile; gradient: 5% B to 65% B in 20 minutes) and purity was calculated to be about 95%, based on the absorbance measured at 280 nm (**Supp. Fig. 2a**). DOPAL unique retention time and profile in RP-HPLC was compared to other catechols (DOPET, DOPAC, epinephrine, norepinephrine, dopamine, L-DOPA, homovanillic acid), confirming the efficacy of the synthesis protocol in converting the epinephrine into DOPAL and in removing the reaction sub-products (**Supp. Fig. 2b**). Consistently, both the absorbance spectrum and the RP-HPLC profile of the synthetized DOPAL perfectly overlap with DOPAL purchased from Santa Cruz Biotechnology (SCBT; sc-391117) that was used as a reference (**Supp. Fig. 2c-d**). Stocks of 100 mM DOPAL in H_2_O were stored at −80° until used.

### Recombinant αSynuclein purification

Recombinant human αSyn was purified as previously described^60^. Briefly, the αSyn gene was cloned in pET-28a plasmid (Novagen) and expressed in *Escherichia coli* BL21(DE3) strain. Bacteria were grown to an OD_600nm_ of 0.3–0.4 and induced with 0.1 mM isopropyl b-D-1-thiogalactopyranoside (IPTG). After 5 hours, cells were collected by centrifugation and recombinant proteins recovered from the periplasm by osmotic shock.

Subsequently, the periplasmic homogenate was boiled for 15 minutes and the soluble αSyn-containing fraction was subjected to a two-step (35 and 55%) ammonium sulfate precipitation. The pellet was then resuspended, extensively dialyzed against 20 mM Tris–HCl, pH 8.0, loaded into a 6 ml Resource Q column (Amersham Biosciences) and eluted with a 0–500 mM gradient of NaCl. Proteins were then dialyzed against water, lyophilized, and stored at −20°C.

### In vitro DOPAL-induced αSynuclein oligomerization

Recombinant αSyn was resuspended in PBS and concentration was determined by measuring the absorbance at 276 nm and using the ε_276nm_ = 5.8 cm^-1^ mM^-1^. αSyn oligomers were produced by incubating recombinant αSyn to a final concentration of 20 µM with 300 µM DOPAL or dopamine (Sigma-Aldrich) in PBS, in a ratio 1:15 αSyn:catechol which corresponds to 1:1 lysine:catechol. The solutions were incubated for 0-2-4-8-16-24-48 hours shaking at 350 rpm at 37 °C. At each time-point, 30 µl of the reactions (10 µg of αSyn) were collected and the reaction was stopped by the addition of loading buffer. Monomeric αSyn and αSyn oligomers were then resolved by SDS-Page into a gradient 4-20% SDS-Page gel (GenScript) and transferred to polyvinylidenedifluoride (PVDF) membranes (BioRad), through a semi-dry Trans-Blot® Turbo™ Transfer System (BioRad). Membrane was scanned at the LiCor Odissey for the nIRF detection at 800 nm, followed by the immunoblot with the rabbit anti-αSyn MJFR1 (ab138501, Abcam) antibody. For the limited proteolysis assay with Proteinase K (PK), DOPAL-αSyn oligomers were produced by overnight incubation of 20 µM αSyn with 600 µM DOPAL in PBS. Monomeric αSyn and DOPAL-αSyn oligomers were then incubated with 50 nM PK (Invitrogen) in PBS for 10-20-30 minutes, and reactions were stopped by the addition of loading buffer. Proteins were then separated by SDS-Page, followed by staining with Coomassie brilliant blue (0.15% Coomassie Brilliant Blue R, 40% ethanol), followed by destaining with 10% isopropanol, 10% acetic acid. Image analysis was then performed with Fiji software.

### Plasmids for mammalian cell expression

The following constructs for primary neuronal cultures and cell line transfection were generated at NCMIR (UCSD): pCAGGS.αSyn-miniSOG full-length; pCAGGS.αSyn-TimeSTAMP-YFP-miniSOG; pCAGGS.αSyn-miniSOG_1-94_(Fragment A) and pCAGGS.αSyn-miniSOG-Jα_95-140_(Fragment B). The pEGFP-N1.αSyn and the pHT2.αSyn-HaloTag for cell line transfection were generated in Prof. Bubacco’s lab (UNIPD). The pEGFP-N1.αSyn was generated from the pEGFP-N1 empty vector (Novagen) as previously described^27^. The pHT2.αSyn(WT)-HaloTag construct was generated by cloning the αSyn-encoding sequence into the pHT2 vector previously described^61^. The αSyn K96R variant was generated using the Quick-Change II site-directed mutagenesis kit (Stratagene) according to manufacturer’s instructions, using the following primers: FOR: 5’ ATTGGCTTTGTCAGAAAGGACCAGTTGG 3’; REV: 5’ CCAACTGGTCCTTTCTGACAAAGCCAGTG 3’.

### Primary mouse cortical neuron preparation and treatments

C57BL/6LOlaHsd and C57BL/6JRccHsd mice, αSyn-null and wild-type respectively, were purchased from Envigo S.r.l. Mouse genotyping was performed with the Phire™ Tissue Direct PCR Master mix (F-170S, ThermoFisher) using primers previously described (mouse Snca exon 6: (AAGACTATGAGCCTGAAGCCTAAG and AGTGTGAAGCCACAACAATATCC, 266 bp fragment; deletion junction, termed D6Slab17: TTGATAGTTCCACTGTTCTGGC and GTAACAATACAGCAAGAGATAC, 179 bp fragment)^62^. Animals were maintained and experiments were conducted according to the Italian Ministry of Health and the approval by the Ethical Committee of the University of Padova (Protocol Permit #200/2029-PR). Cortical neurons were dissociated by papain from P0 pups as previously reported^27^ and cultured in Neurobasal A medium (Life Technologies) supplemented with 2% v/v B27 Supplements (Life Technologies), 0.5 mM L-glutamine (Life Technologies), 100 U/mL penicillin, and 100 µg/mL streptomycin (Life Technologies) for 12 days prior to imaging and western blot analysis, refreshing half medium every three days. Treatments with 100 µM DOPAL were performed in complete medium for 24 hours.

### Primary rat cortical neuron preparation, transfection, and treatments

Cortical neurons were dissociated by papain from postnatal day 2 (P2) Sprague-Dawley rats as previously reported^34^ and according to the animal procedures approved by the Institutional Animal Care and Use Committee of UC San Diego. Neurons were transfected by electroporation using an Amaxa Nucleofection Device (Lonza) at day-in-vitro 0 (DIV0) and cultured in Neurobasal A medium (Life Technologies) supplemented with 1X B27 Supplements (Life Technologies), 2 mM GlutaMAX (Life Technologies), 20 U/mL penicillin, and 50mg/mL streptomycin (Life Technologies) for 10-15 days prior to imaging, refreshing half medium every couple of days. In the αSyn-split-miniSOG experiments, neurons were co-transduced with lentiviral vectors HIV1 expressing αSyn-miniSOG_1-94_(Fragment A) and αSyn-miniSOG-Jα_95-140_(Fragment B) at DIV 7. Treatments with 100 µM DOPAL, 1 µM BILN-2061 (ACME Synthetic Chemical) and 5 µg/ml Nocodazole (487928, Millipore) were performed in complete medium for the time frame specified in each experiment.

### BE(2)-M17 cell line maintenance, transfection, and treatments

Neuroblastoma-derived BE(2)-M17 cells (ATCC CRL-2267) were cultured in 50% of Dulbecco’s modified Eagle’s medium (DMEM, Life Technologies) and 50% of F-12 Nutrient Mix (Life Technologies), supplemented with 10% v/v FBS and 1% Penicillin/Streptomycin (Life technologies). Before transfection and treatments, cells were maintained in complete medium supplemented with 10 µM retinoic acid (RA, Sigma-Aldrich) for three days. When at 80% of confluency, cells were transiently transfected using Lipofectamine 2000 (Invitrogen) with a DNA(µg):Lipofectamine(µl) ratio of 1:2. Cells were then treated and processed 24-to-72 hours post transfection. DOPAL treatments were performed at the final concentration of 100 µM in OptiMEM, unless indicated otherwise. 100 µM DOPAL and 10 mM aminoguanidine hydrochloride (AMG, 396494, Sigma-Aldrich) co-treatments were performed in OptiMEM overnight.

### Catechols detection by nIRF in cell lysate

Catechol-modification of proteins was assessed in DOPAL-treated cells as described in Mazzulli et al. with slight modification^63^. Untreated BE(2)-M17 cells and DOPAL-treated cells were harvested in RIPA Buffer (Cell Signaling Technology) supplemented with protease inhibitors cocktail (Roche). After lysates centrifugation at 20,000 g at 4°C, the cleared supernatant was collected and the protein content in the soluble fraction was determined using the Pierce® BCA Protein Assay Kit (Thermo Scientific) following the manufacturer’s instructions. The insoluble fraction in the pellet of the cell lysate was resuspended in 1 M NaOH (volumes were adjusted according to the protein quantification in the soluble fraction), followed by incubation at 37°C for 30 minutes, sonication at room temperature for 20 minutes (bath sonicator, 100% power and 80 Hz frequency) and centrifugation at maximum speed at room temperature for 20 minutes. The pellets were then dried at 95 °C and resuspended in 10 µl of water. Both the soluble (40 µg of proteins) and the insoluble (4 µl) fractions were spotted on nitrocellulose membrane (BioRad). The near infrared fluorescence signal was acquired by scanning the membrane at the LiCor Odissey using the 800 nm filter and images were analyzed using the Fiji software.

### Western blot

Primary mouse cortical neurons and BE(2)-M17 cells were harvested in RIPA Buffer (Cell Signaling Technology) supplemented with protease inhibitors cocktail (Roche). Lysates were clarified by centrifugation at 20,000 g at 4 °C. Protein concentration in the cleared supernatant was determined using the Pierce® BCA Protein Assay Kit (Thermo Scientific) following the manufacturer’s instructions and protein samples were loaded on gradient 4-20% Tris-MOPS-SDS gels (GenScript). The resolved proteins were then transferred to PVDF membranes (BioRad), through a semi-dry Trans-Blot® Turbo™ Transfer System (BioRad). PVDF membranes were subsequently blocked in Tris-buffered saline plus 0.1% Tween (TBS-T) and 5% non-fat dry milk for 1 hour at 4°C and then incubated over-night at 4°C with primary antibodies diluted in TBS-T plus 5% non-fat milk. The following primary antibodies were used: mouse anti-α-Tubulin (T6074, Sigma-Aldrich), mouse anti-αSyn 211 (S5566, Sigma-Aldrich), mouse anti-αSyn pSer129 81/A (825702, BioLegend), mouse anti-β-Tubulin III (T8578, Sigma-Aldrich), mouse anti-Syn-1 (610787, BD Transduction Laboratories), rabbit anti-αSyn pSer129 EP1536Y (ab51253, Abcam), rabbit anti-αSyn MJFR1 (ab138501, Abcam), mouse anti-β-Actin (A1978, Sigma-Aldrich), mouse anti-Ubiquitin (P4D1 sc-8017, Santa Cruz Biotech.), rabbit anti-p62 (ab109012, Abcam), rabbit anti-Vinculin (AB6039, Millipore), rabbit anti-ALDH1A1 (GTX123973, GeneTex), rabbit anti-ALDH2 (GTX101429, GeneTex), rabbit anti-TH (AB152, Millipore), rabbit anti-α/β.Synuclein (1280 002, SYSY). After incubation with HRP-conjugated secondary antibodies (goat anti-rabbit-HRP and goat anti-mouse-HRP, Sigma-Aldrich) at room temperature for 1 hour, immunoreactive proteins were visualized using Immobilon® Classico Western HRP Substrate (Millipore) or Immobilon® Forte Western HRP Substrate (Millipore) by Imager CHEMI Premium detector (VWR). The densiometric analysis of the detected bands was performed by using the Fiji software.

### Immunofluorescence and confocal microscopy

Rat and mouse primary cortical neurons and BE(2)-M17 cells were fixed using 4% paraformaldehyde (PFA, Sigma-Aldrich) in PBS pH 7.4 for 20 minutes at room temperature. Cells were permeabilized in PBS-0.3% Triton-X for 5 minutes, followed by a 1-hour saturation step in blocking buffer [1% Bovine serum Albumin (BSA) Fraction V, 2% goat serum -for ICC in neurons- or FBS -for ICC in BE(2)-M17 cells-, 0.1% Triton-X and 50 mM Glycine in PBS]. Incubations with primary and secondary antibodies were performed in working solution (1:5 dilution of the blocking buffer) for 1 hour at room temperature, following both incubations with three washing steps in working solution. The following antibodies were used: mouse anti-pSer129 81/A (825702, BioLegend), mouse anti-Syn-1 (610787, BD), chicken anti-β-Tubulin III (302 306, SYSY), rabbit anti-αSyn pSer129 EP1536Y (ab51253, Abcam), mouse anti-aggregated αSyn SynO2 (847601, BioLegend), rabbit anti-aggregated αSyn MKFR14-6-4-2 (ab209538, Abcam), mouse anti-Bassoon (ab82958, Abcam), rabbit anti-VAMP2 (gifted by Prof.

Montecucco’s lab, UNIPD), rat anti-LAMP1 [1D4B] (ab25245, Abcam), rabbit anti-p62 (ab109012, Abcam), mouse anti-Ubiquitin (P4D1 sc-8017, Santa Cruz Biotech.), goat anti-mouse-Alexa Fluor 488 (A11029, Invitrogen), goat anti-mouse-Alexa Fluor 568 (A11004, Invitrogen), goat anti-rabbit-Alexa Fluor 488 (A11034, Invitrogen), rabbit-Alexa Fluor 568 (A11036, Invitrogen), goat anti-rat-Alexa Fluor 647 (A21247, Invitrogen), goat anti-chicken-Alexa Fluor 647 (A21449, Invitrogen). For the immunolabeling with the anti-pSer129 antibody, an antigen retrieval step was introduced before saturation by incubating cells with citrate buffer (10 mM sodium citrate, 0.05% Tween-20, pH 6) at 90°C for 5 minutes or at 70°C for 10 minutes. For the nuclei staining, cells were incubated with Hoechst 33258 (Invitrogen, 1:2000 dilution in PBS) for 5 minutes. Before coverslips mounting on glass slides, cells were washed three times in PBS and rinsed in distilled water. Confocal immunofluorescence z-stack images were acquired on the Olympus Fluoview 1000 laser scanning confocal microscope using a 60X oil immersion objective with numerical aperture 1.42 or on the Zeiss LSM700 laser scanning confocal microscope using a 63X oil immersion objective. To image the fluorescence of αSyn immunolabeling at the confocal microscope, the laser intensity was set based on the αSyn-null neurons acquire αSyn-specific signal.

### Immunofluorescence image analysis

Confocal fluorescence images were processed and analyzed with the Fiji software (https://imagej.net/Fiji). Z-stacks images were converted to maximum intensity z-projections. For the quantification of pSer129 intensity in BE(2)-M17 cells, the Integrated Density (IntDen) was measured on fixed areas in the cytoplasm of transfected cells, normalizing the signal from the immunostaining with the anti-pSer129 antibody to the miniSOG fluorescence (considered as total αSyn). For primary mouse neuron images, immunostaining with anti-β-Tubulin III was used to specifically identify neuronal cells and processes. β-Tubulin III signal was converted to a binary image (setting a threshold) and used as a mask to measure the neuronal area (converted to µm^2^) for further normalization. For the total (Syn-1 immunolabeling) and aggregated (SynO2 and MJFR1-6-4-2 immunolabeling) αSyn fluorescence intensity, the raw IntDen was normalized to the β-Tubulin III area in the field of view, while levels in the soma were measured as Mean Intensity in the region of interest (ROI). To analyze the αSyn puncta fluorescence intensity in the periphery, a fluorescence threshold was set, and the Analyze Particle tool was used. For the quantification of aggregated αSyn puncta (MJFR14-6-4-2 immunolabeling) clustering, the distance among the x,y coordinates of each puncta centroid was measured by the Nearest Neighbor Distance (Nnd) plugin. For the detection of Bassoon-, αSyn- and VAMP2-positive puncta and the percentage of co-localizing puncta, the single channel images were converted to binary images (setting a threshold) and the ComDet v.0.4.2 plugin was used (4.00 pixels particle size, 20.00 intensity threshold, segmentation of larger particles). For the total Ubiquitin fluorescence intensity, the β-Tubulin III mask was subtracted to the Ubiquitin channel and the raw Integrated Density was normalized to the β-Tubulin III area in the field of view. The Ubiquitin levels in the soma were measured as sum of the Integrated Density values of the somata in the field of view, normalized to the total area of the somata; the peripheral Ubiquitin signal was obtained by subtracting the normalized fluorescence in the cell bodies to the total per field of view. For the LAMP1-positive endolysosomes in the neurites, the β-Tubulin III and the LAMP1 channels were converted to binary images (setting a threshold). The images were rotated to visualize each neurite horizontally, and the length was calculated considering the neurite sprout from the soma as X=0 coordinate. Based on the β-Tubulin III binary image, a ROI along the neurite was defined and the LAMP1-positive structures were identified by the Analyze Particle tool (minimum area 0.1 µm^2^) and annotated in terms of number, area and X,Y coordinates of the centroid. The endolysosomal density in the neurites was determined as number of LAMP1-positive particle / neurite length, whereas the endolysosomal proximity to the soma was calculated as percentage of LAMP1-positive particle with X coordinate < 30 µm in each neurite. Finally, to analyze the LAMP1- and p62-positive particles in the soma, a fluorescence threshold was set, and the Analyze Particle tool was used to quantify the particle number, mean size (µm^2^) and mean Fluorescence Intensity.

### Live neuron time-lapse imaging

For live time-lapse recordings, neurons transfected with the αSyn-TimeSTAMP-YFP-miniSOG construct^32, 64^ received a pulse with 1 µM BILN-2061 inhibitor for 4 hours in complete medium at 37°C. After three washes in HBSS, neurons were incubated with imaging medium (HBSS containing 1X B27 Supplements, 25 mM glucose, 1 mM pyruvate, and 20 mM HEPES) in control conditions or with the addition of DOPAL and/or Nocodazole administration. Neurons were imaged with the inverted Olympus Fluoview 1000 laser scanning confocal microscope using a 40X oil immersion objective lens with numerical aperture 1.3 at 0.2% laser power to minimize photo-toxicity. During the time-lapse recordings, cells were maintained at 37°C in controlled atmosphere. Z-stack images were acquired with 200 µm of confocal aperture and 1µm step size (17-24 steps in average) every 30 minutes for 18 hours. Nine different areas were imaged for each independent experiment. Time-lapse movies were edited using Imaris software package. Fluorescence intensity in the cell body was then quantified using IntDen parameter in the Fiji software, whereas the αSyn-positive synapses and relative fluorescence intensity were identified by a custom-made MatLab script (available upon request to the corresponding authors). Only the new terminals identified in the first 3 hours of the time-lapse were considered, assuming t0 as common starting point. For the quantification of the αSyn half-life in the soma and synapses in Nocodazole-treated neurons, the fluorescence variations over time in each individual soma and synapse were fitted by a one-phase decay and the mean value ± SEM was calculated.

### CLEM of miniSOG-tagged αSynuclein

Neurons transfected with the αSyn-TimeSTAMP-YFP-miniSOG construct received a pulse with 1 µM BILN-2061 inhibitor for 4 hours in complete medium. After three washes in complete medium, neurons were kept in the incubator at 37°C in fresh complete medium with or without 100 µM DOPAL. After 24 hours from the BILN-2061 pulse, neurons were fixed and processed for photooxidation and EM processing (next sections below). In the protein fragment complementation assay^35^, neurons that were previously transduced with the two fragments of the αSyn-split-miniSOG, were treated with 100 µM DOPAL in complete medium for 24 hours in the incubator at 37°C. At the end of the treatment, the growth medium was replaced with imaging medium in both the treated and untreated samples and live images of the reconstituted split-miniSOG complex were acquired using the Leica SPE II inverted confocal microscope. Position coordinates were saved. Cells were then fixed in place, and the photo-oxidation protocol followed.

### Confocal fluorescence imaging and photo-oxidation

For the CLEM experiments, cells were fixed using pre-warmed (37°C) 2.5% (w/v) glutaraldehyde (Electron Microscopy Sciences) in 0.1 M sodium cacodylate buffer, pH 7.4 (Ted Pella Incorporated) for 5 minutes at room temperature and then transferred on ice for 1 hour. Subsequently, cells were rinsed on ice 3-5 times using chilled cacodylate buffer and treated for 30 minutes with a blocking solution (50 mM glycine, 10 mM KCN, and 5 mM aminotriazole in 0.1 M sodium cacodylate buffer, pH 7.4) to reduce nonspecific background precipitation of DAB. Cells were imaged using a Leica SPE II inverted confocal microscope outfitted with a stage chilled to 4°C and then photo-oxidized using a 150W Xenon lamp. Confocal fluorescence and transmitted light images were acquired with minimum exposure to identify transfected cells, with care to avoid sample photo-bleaching. For photo-oxidation, oxygenated DAB (3-3’-diaminobenzidine, Sigma-Aldrich) was dissolved in 0.1 N HCl at a concentration of 5.4 mg/ml and subsequently diluted ten-fold into sodium cacodylate buffer (pH 7.4, with a final buffer concentration of 0.1 M), mixed, and passed through a 0.22 mm syringe filter before use. DAB solutions were freshly prepared on the day of photo-oxidation and placed on ice and protected from light before being added to cells. The cells were then illuminated through a standard FITC filter set (EX470/40, DM510, BA520) for miniSOG photo-oxidation, with 100% intense light from a 150 W xenon lamp. Illumination was stopped as soon as an optically-dense reaction product began to appear in place of the fluorescence, as monitored by transmitted light (typically 3–8 min, depending on the initial fluorescence intensity, the brightness of the illumination, and the optics used). Confocal images were analyzed with the Fiji software.

### Electron microscopy

Multiple areas on a single dish were photo-oxidized as described in the previous section. Subsequently, plates with cells were placed on a bed of ice and washed using ice-cold cacodylate buffer to remove unpolymerized DAB. After washing, cells were post-fixed with 1% osmium tetroxide (Electron Microscopy Sciences) in 0.1 M sodium cacodylate buffer for 30 minutes on ice, then washed with ice-cold cacodylate buffer (5 times, 1 minute each) and rinsed once in ice-cold distilled water. An additional staining step with filtered 2% uranyl acetate (UA) in distilled water was performed by overnight incubation at 4°C. The day after, the UA was removed and washed-out with ice-cold distilled water (3 times, 3 minutes each). The samples were then dehydrated with an ice-cold graded ethanol series (20%, 50%, 70%, 90% and100% twice) for 3 minutes each step and washed once in room temperature anhydrous ethanol. Samples were then infiltrated with Durcupan ACM resin (Electron Microscopy Sciences) using a 1:1 solution of anhydrous ethanol:resin for 30 minutes on a platform with gentle rocking, then with 100% resin overnight with rocking. The next day, the resin was removed from dishes (by decanting and gentle scraping with care to avoid touching cells), replaced with freshly prepared resin (3 times, 30 minutes each with rocking), and polymerized in a vacuum oven at 60°C for 48 hours.

Subsequently, photo-oxidized areas of interest were identified by transmitted light, sawed out using a jeweler’s saw, and mounted on dummy acrylic blocks with cyanoacrylic adhesive. The coverslip was carefully removed, and ultrathin sections (80 nm thick) were cut using a diamond knife (Diatome). Electron micrographs were acquired using a FEI Technai 12 (Spirit) transmission electron microscope operated at 80 kV; micrographs were produced using a Tietz 2k by 2k CCD camera and collected using the SerialEM package. Images were then processed and analyzed using Fiji software. Scale bar were adjusted according to the magnification in each image and ultrastructural elements (synaptic vesicles, ILVs and MVBs) were manually annotated. Synaptic vesicles size was expressed as Feret diameter. For the synaptic vesicles clustering analysis, the distance between each pair of vesicles in each pre-synaptic terminal was determined by a custom-made MatLab script (available upon request to the corresponding authors) using the X,Y coordinates of the centroid of each vesicle. The inter-vesicle distances were then visualized by cumulative frequency (bin width 50 nm).

### Pulse-chase experiments with JF_570_-HaloTag ligand

At 24 hours post-transfection, BE(2)-M17 expressing the αSyn(WT)-HaloTag or the αSyn(K96R)-HaloTag constructs, cells were subjected to a pulse with 3 µM JF_570_-HaloTag ligand^65^, diluted in complete medium for 30 minutes. After extensive washes in HBSS to remove the excess of ligand, cells were incubated with 100 µM DOPAL in OptiMEM. After 48 hours, cell medium was washed out and cells were fixed with 4% PFA for 20 minutes at RT. Samples were then washed with PBS and mounted on glass slides with Gelvatol mounting medium (in-house reagent). Z-stacks of JF_570_-positive cells were acquired with the Olympus Fluoview 1000 laser scanning confocal microscope using a 60X oil immersion objective lens with numerical aperture 1.42. Images were analyzed with the Fiji software and the Analyze Particle plugin was used to study αSyn-HaloTag-positive cytoplasmic inclusions.

Alternatively, for the CLEM experiment, cells were fixed with glutaraldehyde following the protocol described for miniSOG, proceeding with the confocal fluorescence imaging at Leica SPEII, photo-oxidation using a ReAsH filter set (P/N:mCherry-A-L01-ZERO, Ex:FF01-562/40(542-582), DM:FF593-Di02, Em:FF01-641/75 (604-679)) and sample processing for EM.

### Pulse-chase experiment with biotin-HaloTag ligand

At 36 hours post-transfection, BE(2)-M17 cells expressing αSyn(WT)-HaloTag or the αSyn(K96R)-HaloTag constructs were treated overnight with 100 µM DOPAL in OptiMEM. At the end of the treatment, cells were subjected to a pulse with 5 µM biotin-HaloTag ligand (G828A, Promega) diluted in complete medium for 3 hours. After three washes in HBSS to remove the excess of ligand, cells were incubated with complete growth medium. At different time-points (t=0-4-8-12-24-30 hours), cells were collected, spinned down and the cell pellets were stored at −80°C. At the end of the time course, all samples were thawed, and cell were harvested in lysis buffer (20 mM Tris-HCl pH7.5, 150 mM NaCl, 1 mM EDTA, 2.5 mM sodium pyrophosphate, 1 mM β-glycerophosphate, 1 mM sodium orthovanadate, 1% Triton® X-100), supplemented with a protease inhibitor cocktail (Sigma-Aldrich). After lysates clarification by centrifugation at 20,000 g at 4°C, protein concentration was determined with the Pierce® BCA Protein Assay Kit (Thermo Scientific) following the manufacturer’s instructions. 150 μg of proteins for each sample were diluted in TBS buffer (Tris-HCl 50 mM pH 7.4, NaCl 150 mM) to a final volume of 300 μl and incubated with 10 μl of medium slurry of streptavidin-coated magnetic beads (GE Healthcare) for 2 hours at room temperature under continuous rotation. The unbounded proteins were then separated by the isolated biotin-labeled αSyn-HaloTag by a magnetic rack and the flow-through was collected separately. The beads were then washed three times with TBS-Urea 2 mM, incubated with 10 μl of loading buffer 1X and boiled at 95°C for 10 minutes. Samples were loaded in gradient 4-20% Tri-Glycine-SDS gels (BioRad) and western blot was performed as follows using the anti-αSyn MJFR1 antibody for the immunoblot. In parallel, 15 μl of flow-through were loaded in a 10% polyacrylamide gel and subjected to an SDS-Page and western blot with the anti-Actin antibody as loading control.

### Zebrafish maintenance and treatment

Wild-type zebrafish were staged and maintained according to standard procedures^66^. Embryos were obtained by natural mating and raised at 28.5°C in Petri dishes containing fish water [50X: 25 g Instant Ocean (Aquarium systems, SS15-10), 39.25 g CaSO_4_ and 5 g NaHCO_3_ for 1 l] with a photoperiod of 13 hours light / 11 hours dark. All zebrafish experiments were carried out at the Zebrafish Facility of the University of Padova (Italy), in accordance with the European (EU 2010/63 directive) and Italian Legislations, under authorization number 407/2015-PR from the Italian Ministry of Health. Zebrafish larvae were exposed to 0.2-0.35 µM disulfiram (86720, Sigma-Aldrich) from 2 dpf to 6 dpf, refreshing the treatment every 24 hours in the morning. The compound was freshly dissolved in DMSO (also used as control) in 20,000X stocks to avoid vehicle-induced toxicity. At 6 dpf, zebrafish larvae were subjected to behavioral analysis followed by whole-mount immunostaining. Alternatively, larvae were euthanized with 0.3 mg/ml tricaine added in fish water and zebrafish heads were isolated by decapitation. The isolated tissues (about 30 heads each group) were homogenized in RIPA buffer supplemented with PMSF and protease inhibitor cocktails (Roche) in ice. Lysates were clarified by centrifugation and stored at −80 °C until western blot analysis as described above, using the mouse anti-βActin, mouse anti-Ubiquitin and rabbit anti-α/β-Synuclein as primary antibodies.

### Swimming behavioral analysis

At 6 dpf, zebrafish larvae were transferred to a 48-wells plate (Sarstedt) 1 hour prior to behavioral analysis. Swimming performance was studied using a DanioVision™ automated tracking system from Noldus Information Technology (Wageningen, the Netherlands) during the light-dark routine according to Lulla et al.^67^, with slight modification. Briefly, larvae were placed inside the Danio Vision Observation chamber (maintained at 28 °C) for 30 minutes in the dark for habituation. The swimming activity was then assessed during three cycles of alternating 10-minutes light and dark periods. The acquired track files were analyzed using the Ethovision X.T. 8.5.614 software (Noldus Information Technology,Wageningen, the Netherlands) for the total distance moved on periods of 2 minutes. Also, the delta distance moved during the light-to-dark stimuli and the total distance moved over the 60 minutes of acquisition were compared among treatments. The experiment was repeated 7 times, using n=8-16 larvae per group for each biological replicate. For the autocorrelation analysis, about 15 larvae per conditions form three independent experiments were analyzed. The velocity tracks (10 second-bin) of the movement acquired during the light-dark routine test was analyzed by Simple Time Series Analysis plugin of OriginPro 2020 with Bartlett’s approximation and plotted as AutoCorrelation Function (ACF) ± SEM.

### Zebrafish whole-mount immunofluorescence

After swimming behavioral analysis, zebrafish larvae were euthanized with 0.3 mg/ml tricaine and fixed overnight in 4% PFA (Sigma) in PBS while still in the 48-wells plate. Larvae were then post-fixed in 100% methanol and separately transferred to 1.5 ml-tubes keeping track of their position on the plate of the behavioral analysis. After rehydration with graded methanol series (75-50-25% in PBS) and wash in 0.2% Triton in PBS (PBT) for 5 minutes at room temperature, larvae were incubated in 3% H_2_O_2_ - 5% KOH in water for 20 minutes at room temperature for depigmentation. Larvae were then permeabilized in ice-cold 100% acetone for 15 minutes at −20 °C, following washes in distilled water and PBT. After a saturation step in 1% BSA, 2% goat serum in PBT (PBTB) for 4 hours at room temperature, larvae where incubated with rabbit anti-TH primary antibody (AB152, Millipore) diluted 1:100 in PBTB for 24 hours at 4 °C. The following day, larvae were washed 4 times in PBT for 20 minutes each at RT prior to overnight incubation with goat anti-rabbit Alexa Fluor 488 (A11034, Invitrogen) diluted 1:100 in PBTB at 4 °C in the dark. After extensive washes in PBT, larvae were embedded in 0.8% low-melting agarose and placed on depression slides. TH-positive neuronal clusters in the brain were imaged by z-stacks (about 25 steps, 10 µm step size) under a 10X dry objective at the Zeiss LSM700 laser scanning confocal microscope. The acquired images were then analyzed with Fiji software. The fluorescence intensity (maximum intensity z-projections) of the TH-positive neuronal clusters in the DE and the OB&TE was measured as IntDen on a fixed area among samples, subtracting the background signal to each image.

### Synuclein sequences alignment

Protein sequences of mammalian αSyn (P37840_Human, O55042_Mouse, P37377_Rat) and zebrafish Synuclein isoforms (Q7SX92_Sncb, Q568R9_Sncga, Q502J6_Sncgb) were obtained from UniProtKB (https://www.uniprot.org). Multiple sequence alignment was performed by CLUSTALW (https://www.genome.jp/tools-bin/clustalw) and visualized by Jalview software (https://www.jalview.org).

### *Aldh1a1^-/-^/Aldh2^-/-^* mice and brain tissue preparation

Mice with homozygous deletions of both *Aldh1a1* and *Aldh2* genes (double knock-out mice) on a C57BL/6 background were generated and genotyped as previously described^43^. Animals were maintained and experiments were conducted according to the National Institute of Health “Guide for the Care and Use of Laboratory Animals” and the approval by the Institutional Animal Care and Use Committee of the University of Texas Health Science Center at San Antonio. Age-matched male *Aldh1a1^-/-^/Aldh2^-/-^* and wild-type mice were sacrificed between 11 and 12 months of age. Brains were rapidly collected following brief carbon dioxide anesthesia and decapitation. For western blot assays, brain tissues were rapidly dissected on an ice-cold glass plate, collecting the midbrain and striatum separately; tissues were then snap-frozen on dry ice and transferred to a –80°C freezer for storage until assayed. Samples were homogenized in freshly prepared ice-cold RIPA buffer (Cell Signaling Technology) and supplemented with phosphatase inhibitors cocktail (Life Technologies) and protease inhibitors (Roche) cocktails. Clarified lysates were then analyzed by SDS-Page and immunoblotting.

## Statistical analysis

The quantitative and statistical analysis of the collected data, as well as the proper graphical visualization, were performed by the GraphPad Prism Software Inc. (version 7). In general, data were collected from at least n=3 independent experiments, with multiple technical replicates for each group. Within each biological replicate, data were normalized to the mean value of the untreated condition and plotted as fold-change variations of the treated samples compared to controls. Data from independent experiments were then pooled together and represented as box plots showing median, 25^th^ and 75^th^ percentiles, minimum and maximum values. The statistical analysis among mean values was performed by two-tailed non-parametric Mann-Whitney test (two datasets) or by the two-tailed non-parametric Kruskall-Wallis test with the Dunn’s multiple comparisons test (more than two datasets). For grouped analysis, the two-way ANOVA test with the Sidak’s multiple comparison test was performed. Statistical significance was defined for p-value < 0.05 (ns p>0.05, * p<0.05, ** p<0.01, *** p<0.001, **** p<0.0001). Additional details on the data analysis are reported in the legend of each figure.

---

**Supporting Figure 1.**
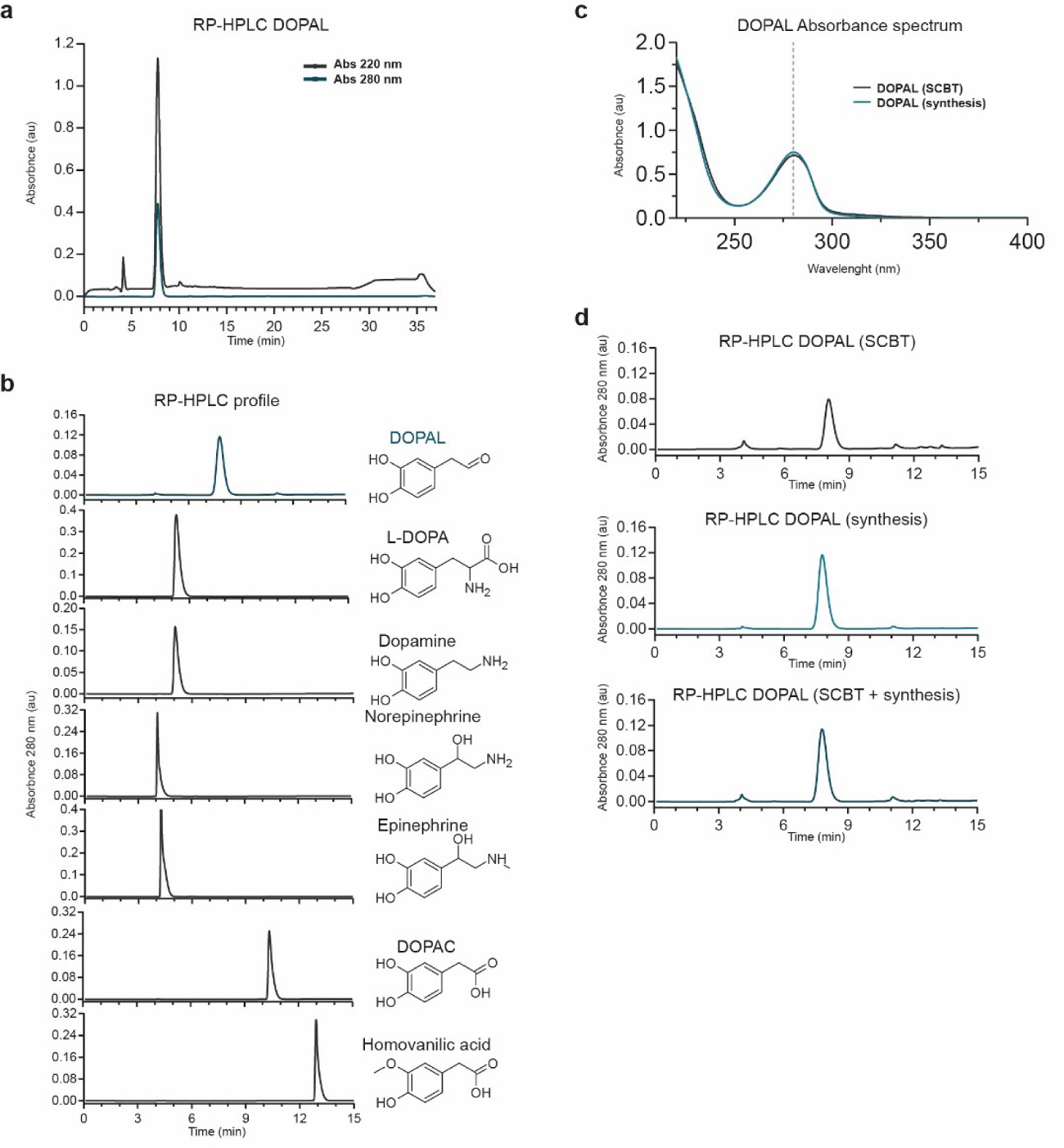
DOPAL quality control analysis. (**a**) RP-HPLC of the synthetized DOPAL with the absorbance profiles at 220 nm (in grey) and 280 nm (in green). (**b**) Comparison of the RP-HPLC retention time and profile (absorbance at 280 nm) among the synthetized DOPAL and other catechols. The chemical structures of the different molecules are indicated on the right side of the corresponding RP-HPLC profile. (**c**) Overlap between the absorbance spectra of the synthetized DOPAL (in green) and the DOPAL purchased from Santa Cruz BioTech. (SCBT, in grey). The dotted line indicates the absorbance peak at 280 nm. (**d**) Comparison of the RP-HPLC retention time and profile among the synthetized DOPAL, the DOPAL by SCBT and the combination of the two.

**Supporting Figure 2.**
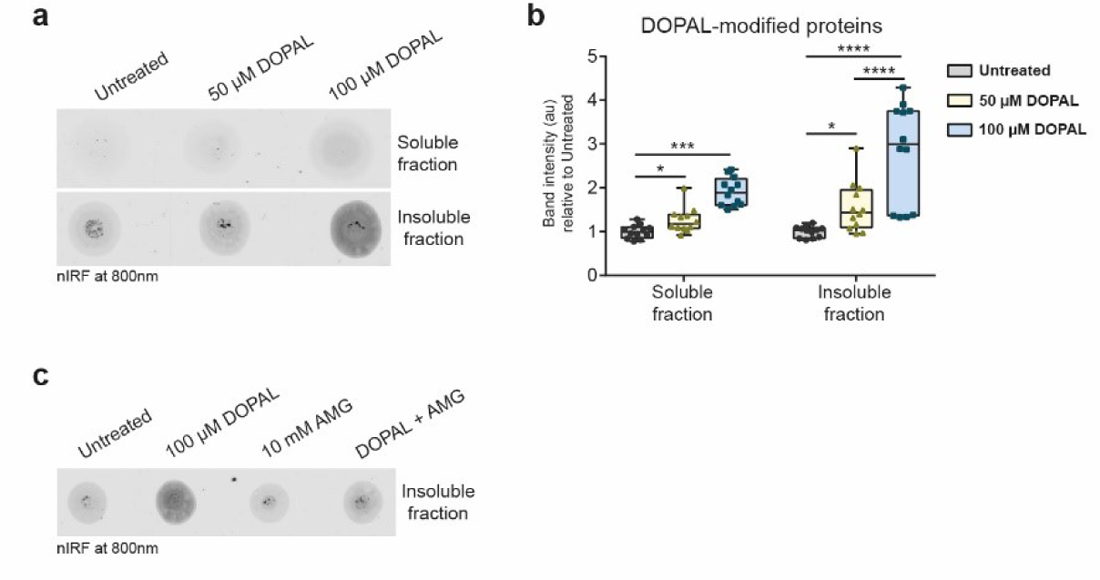
DOPAL cellular treatment results in a dose-dependent accumulation of DOPAL-modified proteins. (**a**) DOPAL-derived nIRF signal in detergent-soluble and insoluble fractions of BE(2)-M17 after an overnight treatment with 0-50-100 µM DOPAL, and (**b**) corresponding quantification. Data from three independent experiments (with n=4 technical replicates each, pooled together) are normalized to each untreated sample and analyzed by Two-way ANOVA with Sidak’s multiple comparison test (* p<0.05, *** p<0.001, **** p<0.0001). (**c**) DOPAL-derived nIRF signal in detergent-insoluble fraction of BE(2)-M17 after an overnight treatment with 100 µM DOPAL or the co-treatment with 10 mM AMG.

**Supporting Figure 3.**
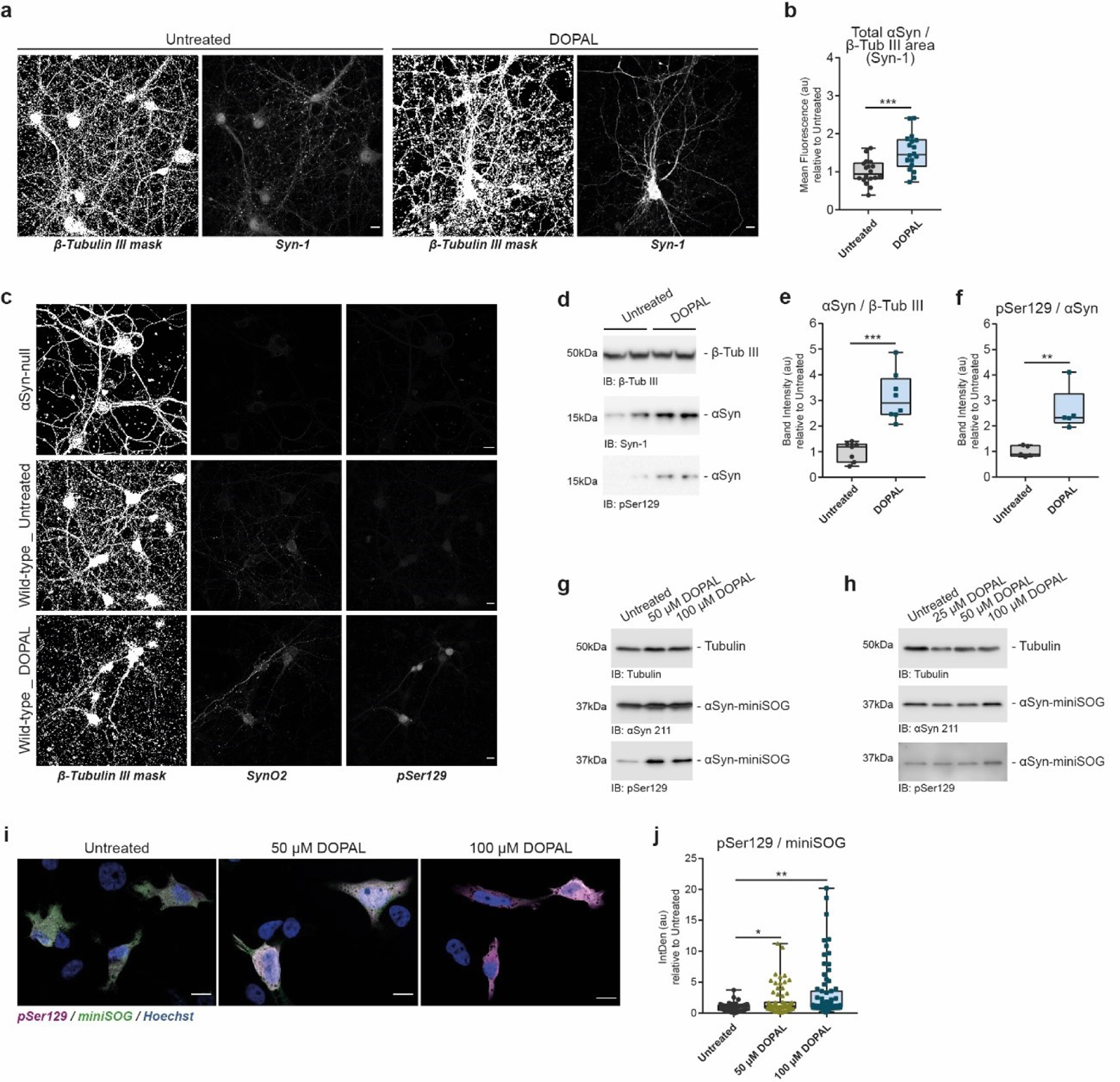
DOPAL induces αSynuclein accumulation and increased αSynuclein phosphorylation at Serine 129. (**a**) Immunostaining of β-Tubulin III (converted to binary mask) and total αSyn (Syn-1) in untreated and 100 μM DOPAL-treated (for 24 hours) wild-type primary mouse cortical (referred to Fig. 1b). Scale bar: 10 μm. (**b**) DOPAL-induced αSyn accumulation is expressed as total αSyn fluorescence normalized to β-Tubulin III-positive area. Data from three independent cultures are normalized to each untreated sample, pooled together, and analyzed by Mann-Whitney non-parametric test (*** p<0.001). (**c**) Immunostaining of β-Tubulin III (converted to binary mask), oligomeric (SynO2) and phosphorylated (pSer129) αSyn in untreated αSyn-null and wild-type, and 100 μM DOPAL-treated (for 24 hours) wild-type primary mouse cortical neurons. (**d-e**) Total αSyn and (**f**) pSer129 levels in untreated and 100 μM DOPAL-treated (for 24 hours) wild-type primary mouse cortical neurons, analyzed by western blot. Band intensities are normalized to β-Tubulin III as loading control. Data from three independent cultures are normalized to each untreated sample, pooled together, and analyzed by Mann-Whitney non-parametric test (** p<0.01, *** p<0.001). DOPAL dose-dependent increase of αSyn phosphorylation at Ser129 analyzed by western blot in (**g**) BE(2)-M17 cells and (**h**) primary rat cortical neurons overexpressing αSyn-miniSOG after an overnight treatment with 0-25-50-100 µM DOPAL. In the same experimental condition, (**i**) the increased pSer129 was confirmed in BE(2)-M17 cells overexpressing αSyn-miniSOG by immunofluorescence. The fluorescence intensity of the immunolabeling with the anti-pSer129 antibody normalized to the miniSOG fluorescence in each cell. Nuclei are stained with Hoechst (blue). Scale bar: 10 µm. (**j**) Data from three independent experiments are normalized to each untreated sample, pooled together, and analyzed by Kruskall-Wallis non-parametric test with Dunn’s multiple comparison test (* p<0.05, ** p<0.01).

**Supporting Figure 4.**
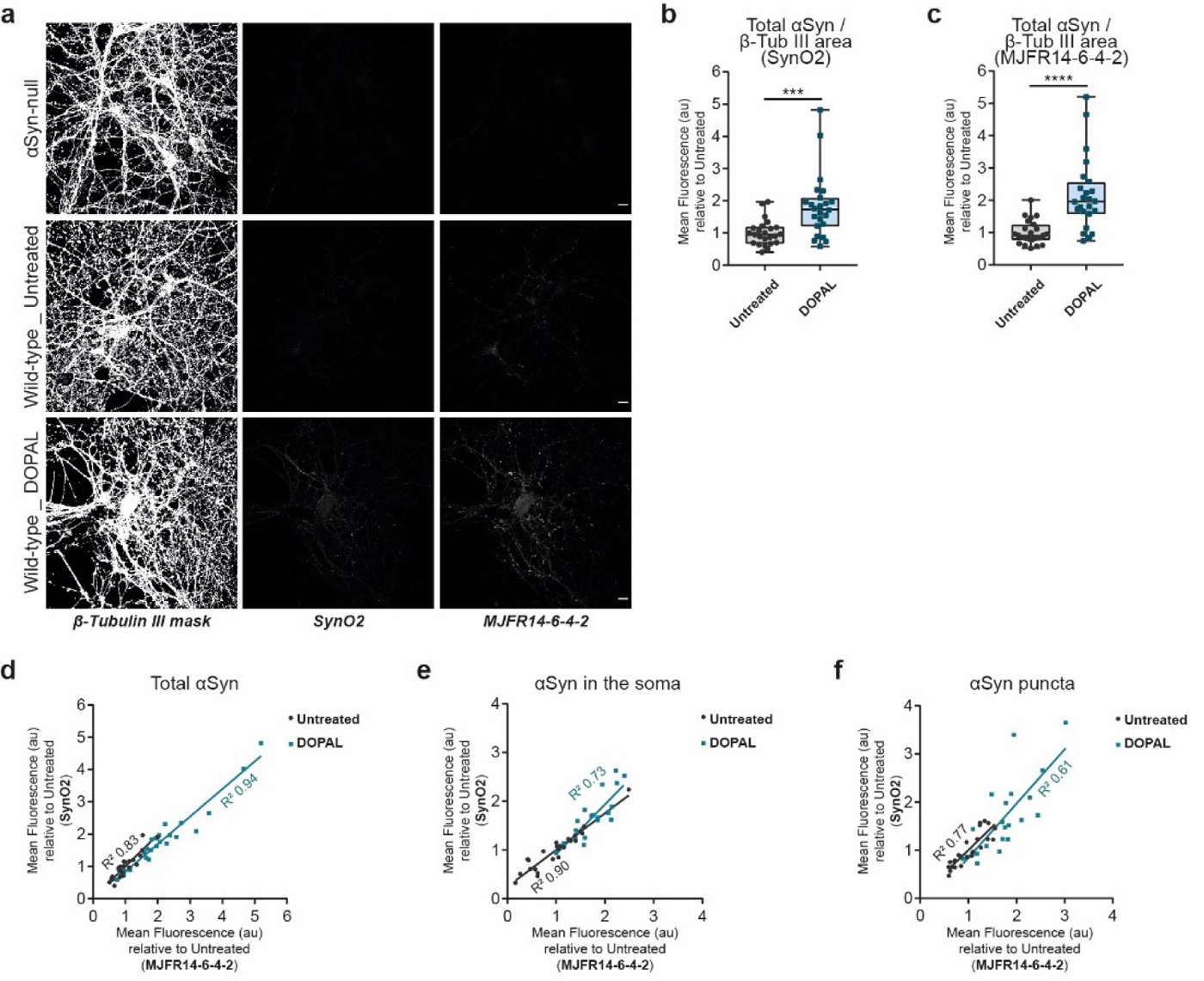
DOPAL induces multimeric αSynuclein buildup in primary mouse neurons. (**a**) Immunostaining of β-Tubulin III (converted to binary mask), oligomeric (SynO2) and aggregated (MJFR14-6-4-2) αSyn in untreated αSyn-null and wild-type, and 100 μM DOPAL-treated (for 24 hours) wild-type primary mouse cortical neurons (referred to Fig. 1f). Scale bar: 10 µm. (**b-c**) DOPAL-induced multimeric αSyn accumulation is expressed as αSyn fluorescence normalized to β-Tubulin III-positive area for both immunostainings with the two antibodies. Data from two independent cultures are normalized to each untreated sample, pooled together, and analyzed by Mann-Whitney non-parametric test (*** p<0.001, **** p<0.0001). Linear correlation between the DOPAL-induced increased fluorescence intensities derived from the immunostaining with SynO2 and MJFR14-6-4-2 antibodies, measured (**d**) in the entire field of view (total), (**e**) in the soma and (**f**) in the synaptic puncta.

**Supporting Figure 5.**
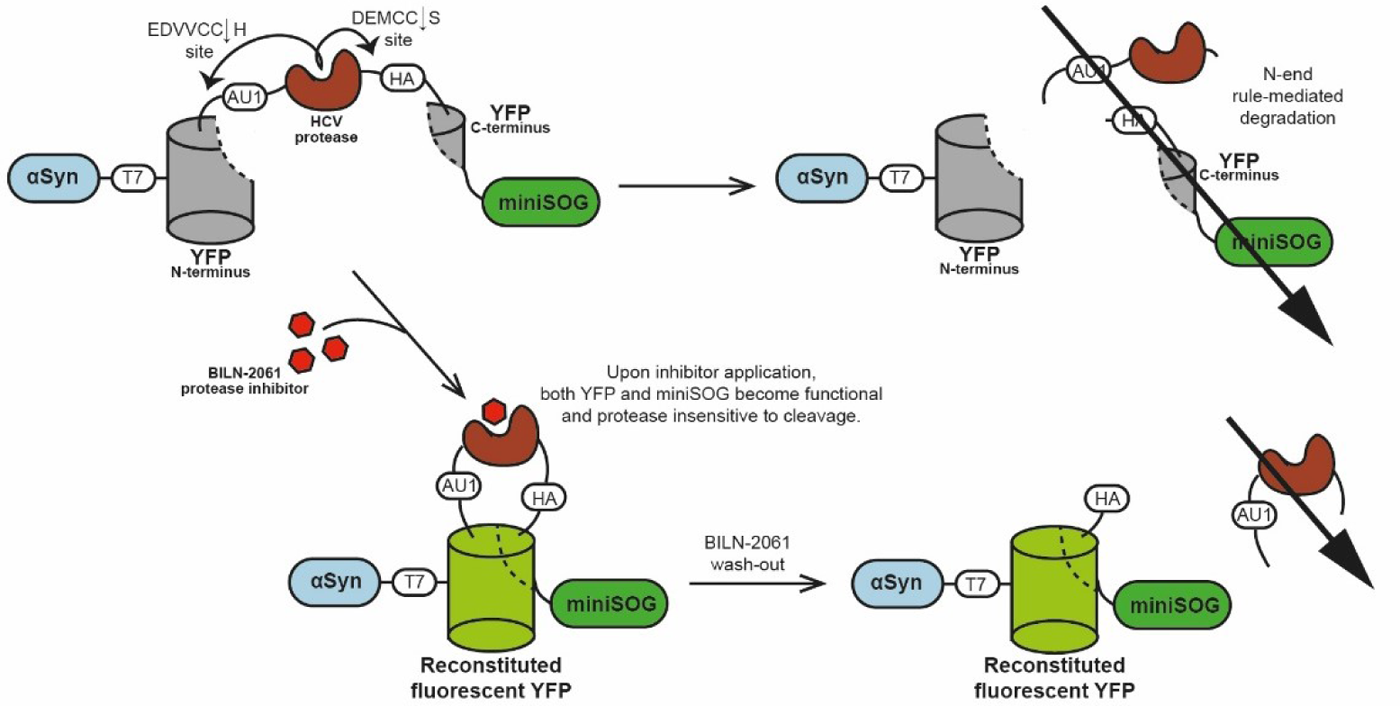
Schematic representation of αSynuclein-TimeSTAMP-YFP-miniSOG. TimeSTAMP (Time-Specific Tag for the Measurement of the Age of Proteins) is a drug-sensitive on-off switch with a cassette comprising the protease domain of hepatitis C virus (HCV) flanked by protease cleavage sites and followed by a split-YFP epitope tag. Upon expression, the protease excises itself and the tag from proteins by default, targeting the two fragments to degradation. However, by applying a cell-permeant protease inhibitor (BILN-2061), αSyn is expressed in frame with a reconstituted and active YFP probe, providing the capability to select and image by LM a newly synthetized protein subpopulation. Here, the TimeSTAMP tag was combined with YFP fluorescent protein and miniSOG, to analyze αSyn trafficking by both live-imaging and CLEM. *Adapted from Butko et al., Nat Neurosci 2012*.

**Supporting Figure 6.**
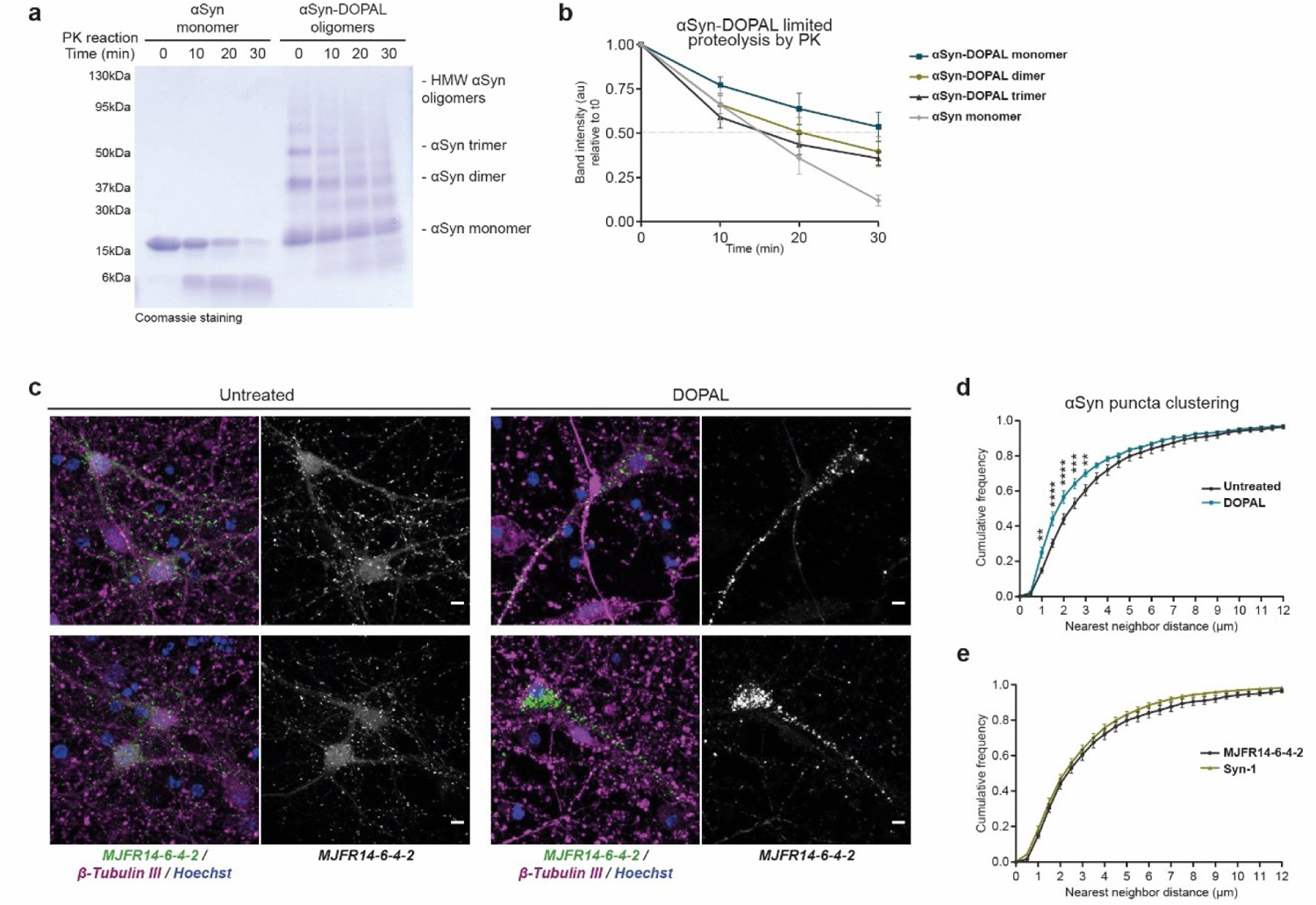
DOPAL affects αSynuclein proteolysis *in vitro* and induces aggregated αSynuclein accumulation in primary neurons. (**a**) SDS-Page and Coomassie staining of the *in vitro* limited proteolysis assay by PK (0-10-20-30 minutes time-course) of 20 µM recombinant monomeric αSyn and αSyn-DOPAL oligomers obtained by the incubation of 20 µM αSyn with 300 µM DOPAL overnight. (**b**) In the quantification, the levels of αSyn monomers, dimers and trimers over time are normalized to the amount at t0. Data are presented as mean ± SEM from four independent experiments. (**c**) Immunostaining of β-Tubulin III (magenta) and aggregated αSyn (MJFR14-6-4-2, green) in untreated and 100 μM DOPAL-treated (for 48 hours) wild-type primary mouse cortical neurons. Nuclei are stained with Hoechst (blue). Scale bar: 5 µm. (**d**) The clustering of aggregated αSyn-positive puncta is expressed cumulative frequency of the nearest neighbor distance (µm) between puncta centroids. Data are pooled from three independent cultures and analyzed by Two-way ANOVA with Sidak’s multiple comparison test (** p<0.01, *** p<0.001, **** p<0.0001). (**e**) Conversely the distance distribution among αSyn-positive puncta detected by Syn-1 staining (images refer to Fig. 1b) doesn’t differ from the puncta detected by the MJFR14-6-4-2 antibody, both analyzed in untreated neurons.

**Supporting Figure 7.**
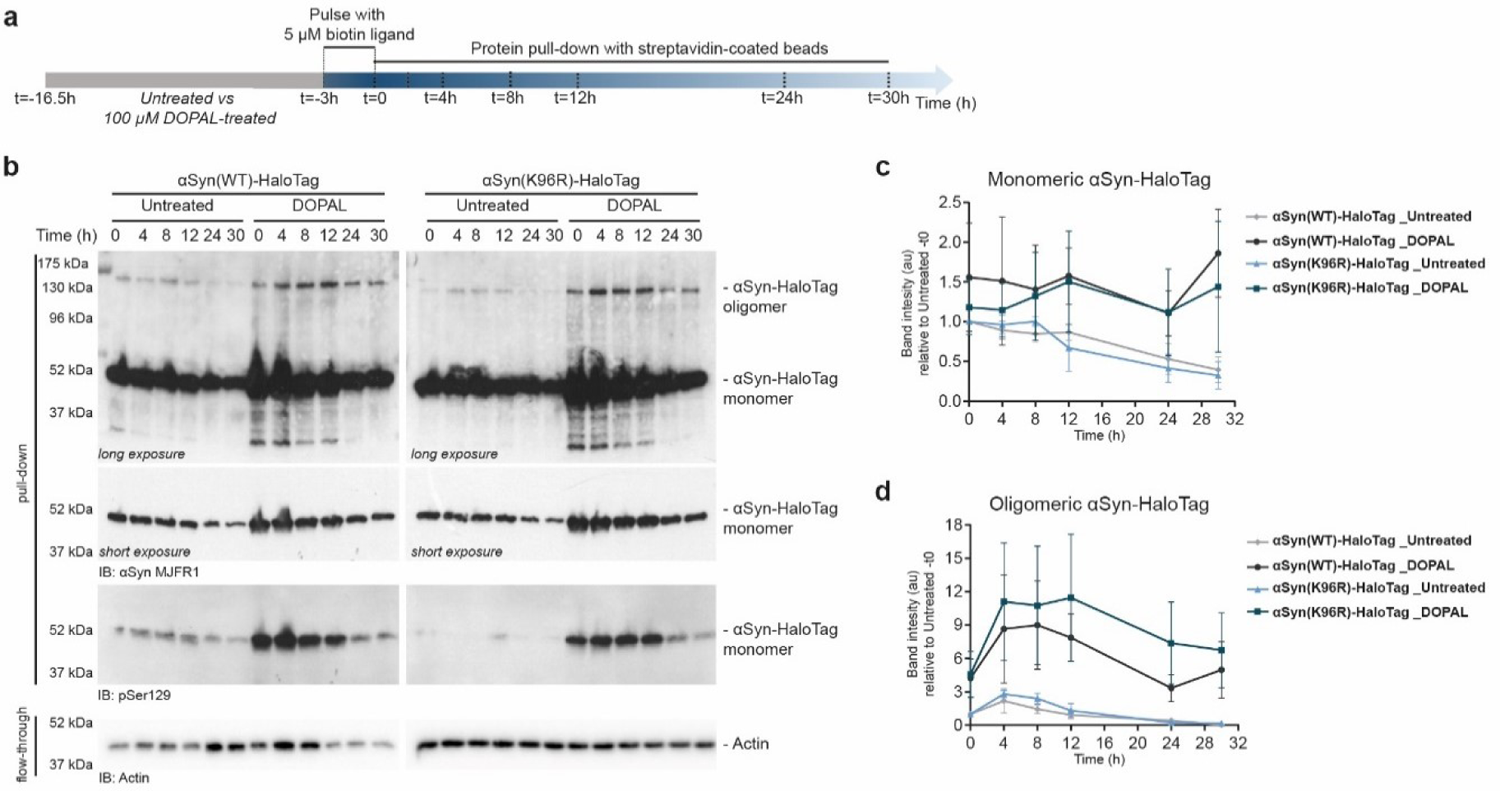
DOPAL affects the turn-over of WT and K96R αSynuclein in BE(2)-M17 cells. (**a**) Schematic representation of the pulse-chase experiment using the biotin HaloTag ligand in αSyn-HaloTag overexpressing BE(2)-M17 cells. (**b**) Western blot of the degradation of monomeric and oligomeric αSyn(WT)-HaloTag and αSyn(K96R)-HaloTag, without and with 100 μM DOPAL treatment. (**c-d**) In the quantification, the αSyn band intensity in the pull-down was normalized to the actin in the flow-through and to the starting amount at t0 in the untreated cells for each αSyn variant. Data are presented as mean ± SEM from n=3 independent experiments and analyzed by Two-way ANOVA: (**c**) αSyn(WT)_ interaction p=0.24, time p=0.33, treatment p=0.22; αSyn(K96R)_ interaction p=0.19, time p=0.43, treatment p=0.35; (**d**) αSyn(WT)_ interaction p=0.42, time p=0.06, treatment p=0.10; αSyn(K96R)_ interaction p=0.14, time p=0.003, treatment ns p=0.16.

**Supporting Figure 8.**
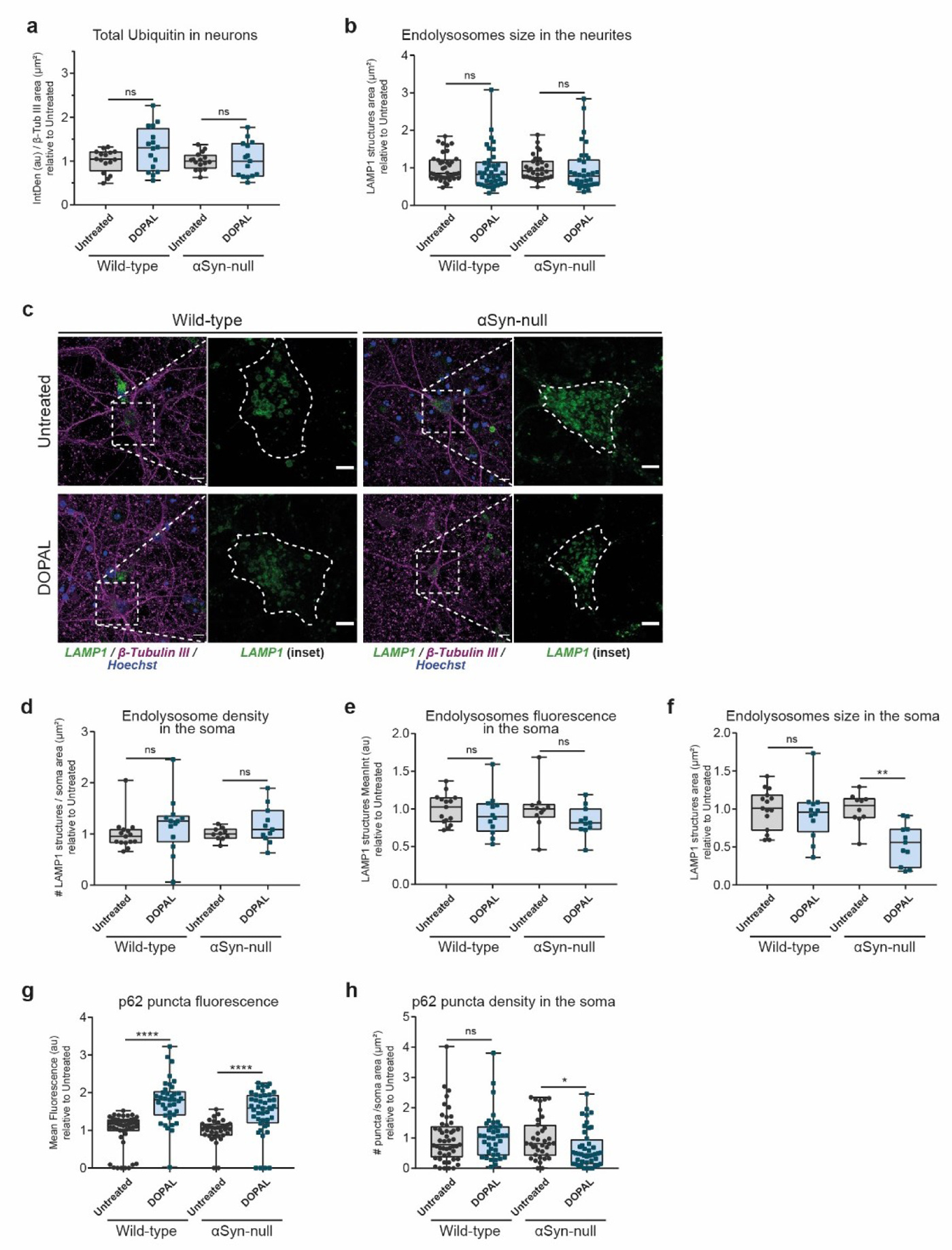
Additional quantification of Ubiquitin, LAMP1 and p62 in primary mouse neurons. (**a**) Quantification of the total Ubiquitin fluorescence signal in neurons, normalized to β-Tubulin III area (referred to Fig. 4a). (**b**) Quantification of the LAMP1 structures mean size (µm^2^) in each neurite (referred to Fig. 4d). (**c**) Immunostaining of β-Tubulin III (magenta) and LAMP1 (green) in the soma of untreated and 100 μM DOPAL-treated (for 24 hours) wild-type and αSyn-null primary mouse cortical neurons. Nuclei are stained with Hoechst (blue). Scale bar: 10 µm. In the inset, the LAMP1 fluorescence signal in the cell body is enlarged and the dotted line defines the soma boundaries (scale bar: 5 µm). Quantification of (**d**) endolysosomes density in the cell body expressed as number of LAMP1 structures / µm^2^, (**e**) LAMP1 structures mean fluorescence signal and (**f**) LAMP1 structures mean size (µm^2^) in each cell body. Quantification of (**g**) p62-positive puncta mean fluorescence signal and (**h**) p62-positive puncta mean size (µm^2^) in each cell body (referred to Fig. 4g). (**a, b, d, e, f, g, h**) Data from three independent experiments are normalized to each untreated sample, pooled together, and analyzed by Mann-Whitney non-parametric test (* p<0.05, ** p<0.01, **** p<0.0001).

**Supporting Figure 9.**
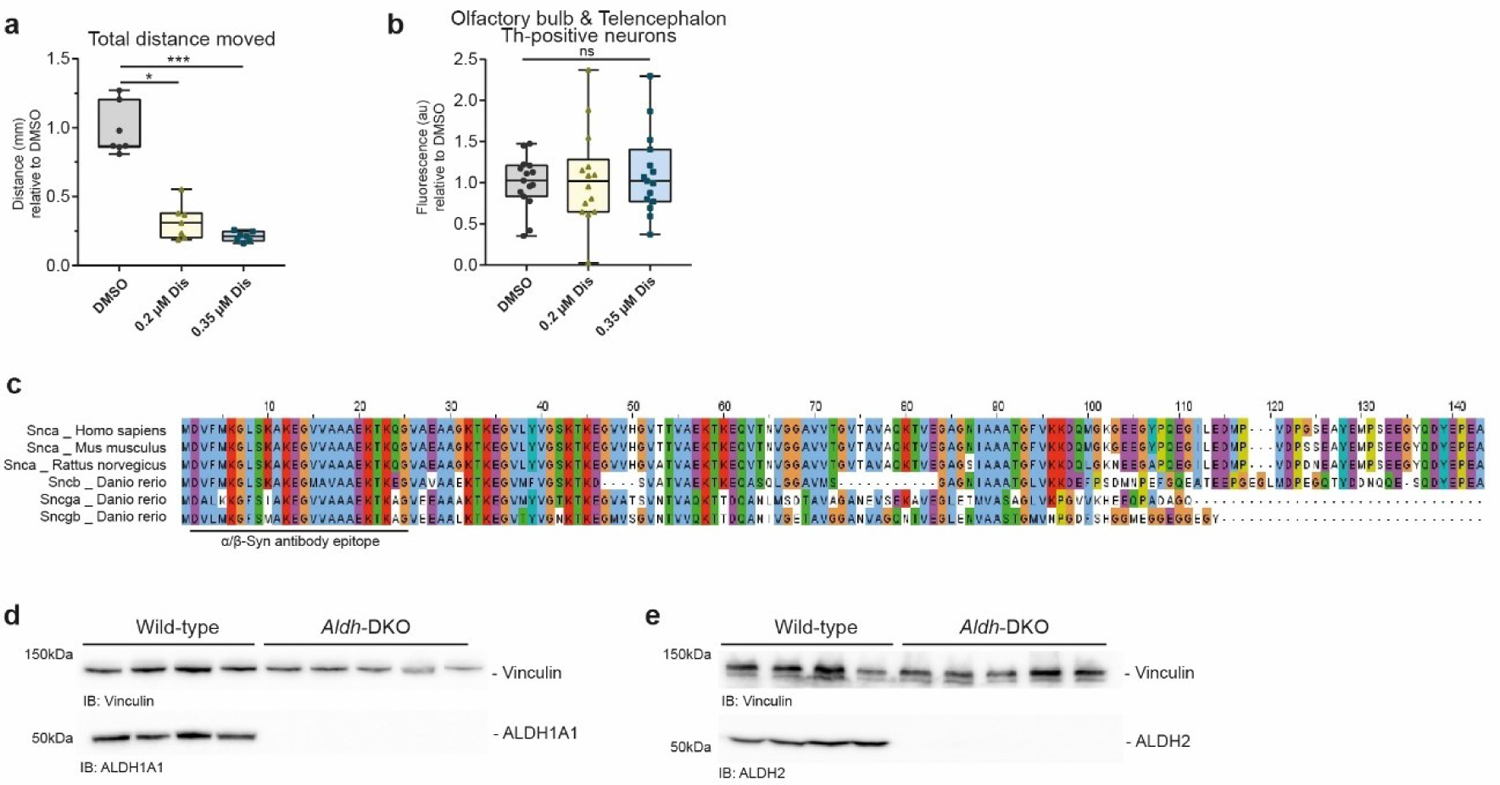
Additional data on *in vivo* models. (**h**) Quantification of the total distance moved during the light-dark locomotion test of DMSO-, 0.2 µM Disulfiram and 0.35 µM Disulfiram-treated zebrafish larvae (referred to Fig. 5a). Data from seven independent experiments are analyzed by two-way ANOVA with Tukey’s multiple comparison test (** p<0.01, **** p<0.0001). (**b**) Quantification of TH fluorescence signal in the dopaminergic neuron cluster in the olfactory bulb & telencephalon (referred to Fig. 5d). Data from three independent experiments are normalized to each DMSO sample, pooled together, and analyzed by Kruskall-Wallis non-parametric test with Dunn’s multiple comparison test. (**c**) Aminoacidic sequence alignment of Snca protein of *Homo sapiens*, *Mus musculus* and *Rattus norvegicus* with the Sncb, Sncga and Sncgb of *Danio rerio* obtained by CLUSTALW and visualized by Jalview software. The sequence epitope 2-25 aa of the anti α/β-Syn antibody (SYSY) is reported. Western blot of wild-type (n=4) and ALDH-DKO (n=5) mice (**d**) *striatum* and (**e**) cortex with the antibodies against ALDH1A1 and ALDH2. Immunoblot against Vinculin was used as loading control.

**Supporting Figure 10.**
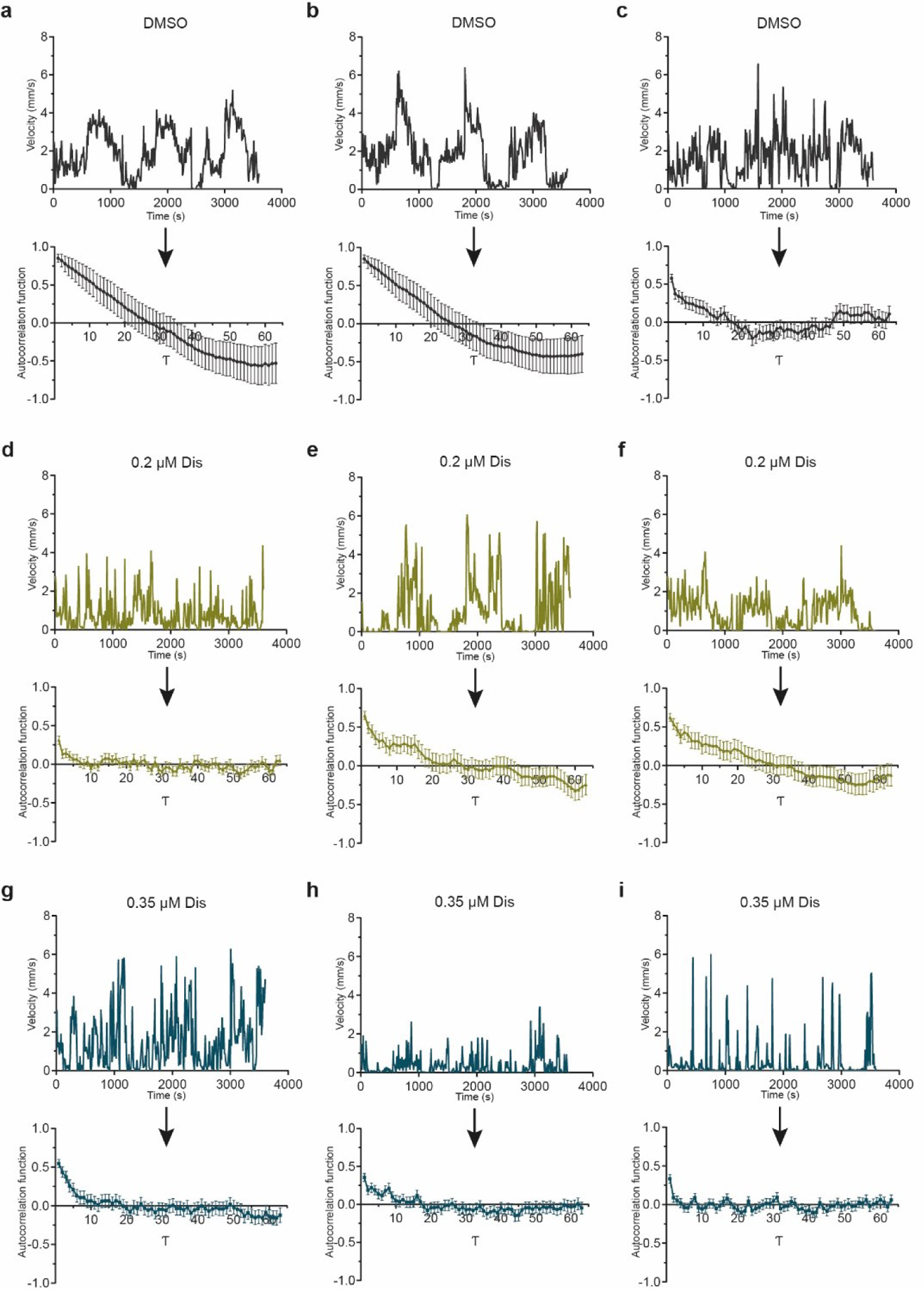
Examples of autocorrelation analysis of zebrafish swimming behavior. Representative graphs showing the time-course of velocity (10 second-bin) of movement during the light-dark routine test acquired at DanioVision™ and the corresponding autocorrelation function of zebrafish larvae exposed to (**a-c**) DMSO, (**d-f**) 0.2 µM Disulfiram and (**g-i**) 0.35 µM Disulfiram.

